# Structural basis and functional roles for Toll-like receptor binding to Latrophilin adhesion-GPCR in embryo development

**DOI:** 10.1101/2023.05.04.539414

**Authors:** Gabriel Carmona-Rosas, Jingxian Li, Jayson J. Smith, Shouqiang Cheng, Elana Baltrusaitis, Wioletta I. Nawrocka, Minglei Zhao, Paschalis Kratsios, Demet Araç, Engin Özkan

## Abstract

Latrophilins/ADGRLs are conserved adhesion-type G protein-coupled receptors associated with early embryonic morphogenesis defects, lethality, and sterility across multiple model organisms. However, their mechanistic roles in embryogenesis and the identity of their binding ligands remain unknown. Here, we identified a cell-surface receptor, TOL-1, the sole Toll-like receptor in *C. elegans*, as a novel ligand for the *C. elegans* Latrophilin, LAT-1. The extracellular lectin domain of LAT-1 directly binds to the second leucine-rich repeat domain of TOL-1. The highresolution crystal structure and the cryo-EM density map of the LAT-1–TOL-1 ectodomain complex reveal a previously-unobserved mode of one-to-one interaction enabled by a large interface. CRISPR/Cas9-mediated mutation of key interface residues selectively disrupted the endogenous LAT-1–TOL-1 interaction in *C. elegans,* leading to partial sterility, lethality, and malformed embryos. Thus, TOL-1 binding to LAT-1 represents a receptor-ligand axis essential for animal morphogenesis.

## INTRODUCTION

During animal development, cells communicate with each other to follow a sequence of precisely controlled cell division events and to spatially arrange in a coordinated manner forming the embryo. Cellular communication during this process is mediated by cell-surface receptors that are believed to be conserved across animals. Several key cell-surface receptors mediate extracellular protein-protein interactions to induce signaling pathways acting at different times and in different regions of the embryo.^1^ However, the molecular mechanisms of cellular communication that govern early embryogenesis are incompletely understood.

Latrophilins (LPHNs/ADGRLs) are cell-surface receptors with essential functions in animal development, especially in early embryogenesis, in both vertebrates and invertebrates.^2, 3^ They are signaling receptors that belong to the adhesion G-protein-coupled receptor (aGPCR) family, and are found in all multicellular animals.^4–7^ Among the 33 aGPCRs in humans, LPHNs (LPHN1-3) are the only aGPCRs, besides flamingo-like Cadherin EGF LAG seven-pass G-type receptors (CELSRs), that are conserved between vertebrates and invertebrates.^8^ LPHNs have large extracellular domains (ECDs) comprised of a Lectin, an Olfactomedin, a Hormone-Binding and a GAIN domain, which precedes the transmembrane (7TM) region (**Fig. 1A**).^9–11^ The olfactomedin domain is absent in the *C. elegans* homolog, LAT-1.

**Figure 1.**
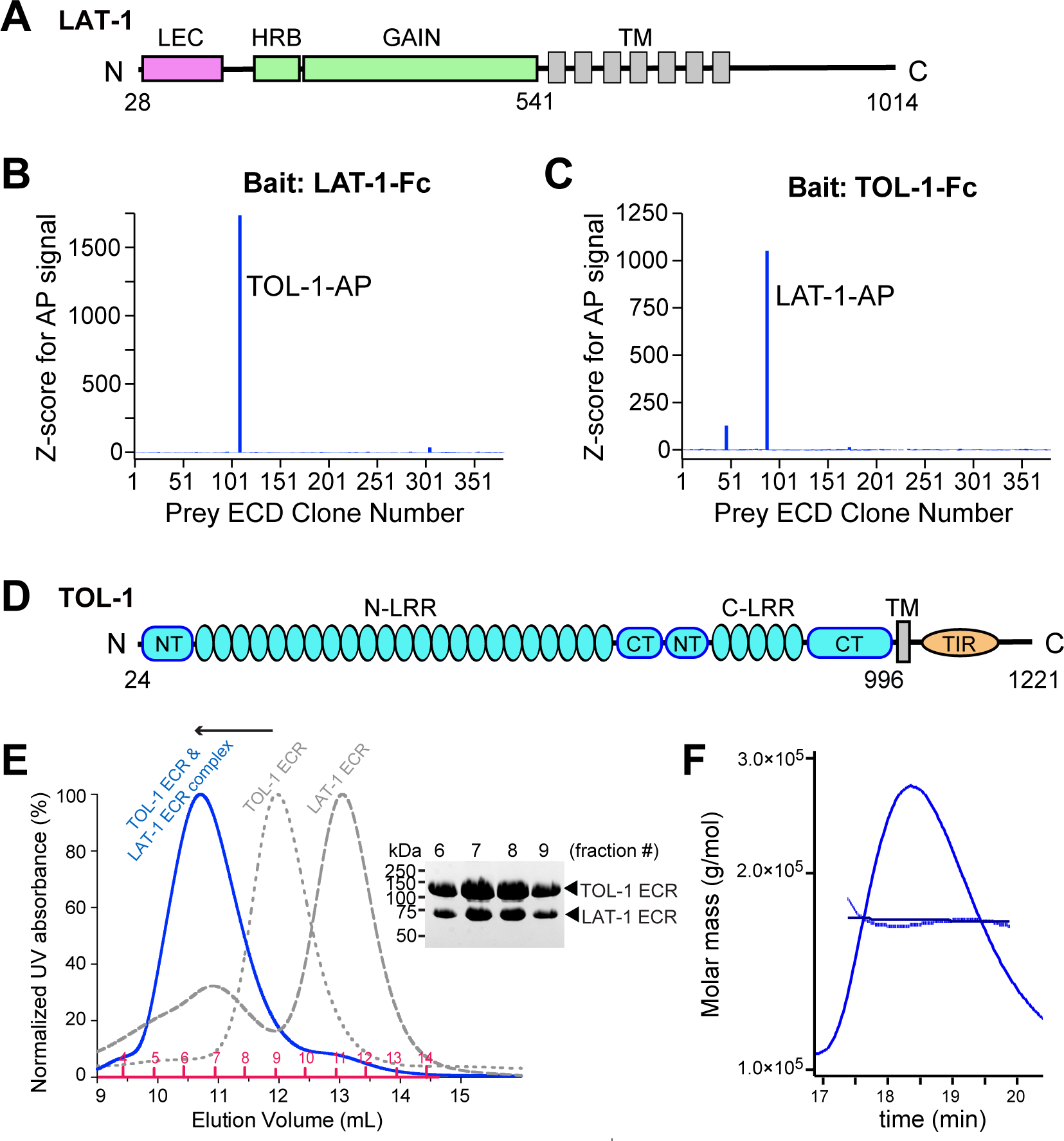
LAT-1 binds to TOL-1 with high affinity (A) Domain architecture schematics of LAT-1, the *C. elegans* homolog of Latrophilin. (C, D) *Z*-scores for AP signal observed against a LAT-1-Fc bait (B) and TOL-1-Fc bait (C). Highest signals are observed for AP-tagged TOL-1 (B) and LAT-1 (C). (B) Domain architecture schematics of TOL-1, the only *C. elegans* homolog of Toll-like receptors. (E) SEC profiles of full ECDs of TOL-1, LAT-1 and their complex; and the SDS-PAGE analysis of the fractions form the complex peak. Numbers indicate the collected fractions. (F) MALS analysis of the complex peak showing a molecular weight of 172 kDa corresponding to a 1:1 stable complex of TOL-1 (112.1 kDa) and LAT-1 (58.9 kDa) which adds up to 171 kDa). Also see Supplementary Fig. 1.

The roles of LPHNs in embryonic development have been mostly studied in *C. elegans*, whose genome contains two LPHN orthologs, *lat-1* and *lat-2*. Although the function of *lat-2* remains unknown, *lat-1* is essential for early development; it is required for the alignment of cell division planes along the anterior-posterior axis.^12^ *lat-1* is expressed in early blastomeres, developing embryonic epithelia, specifically in left epidermoblasts during dorsal intercalation, in pharyngeal muscles and neurons, and in the nerve ring.^12, 13^ Its expression is needed for embryo elongation, pharyngeal development, and sperm development and/or function. *lat-1* null mutant worms display embryonic and larval lethality phenotypes (17% and 56%). Mice carrying homozygous null *Lphn1*^14, 15^ or *Lphn2*^16^ alleles also show embryonic lethality, indicating that LPHN function in embryogenesis is conserved between invertebrates and vertebrates. Finally, numerous recent studies suggest roles for LPHNs in tissue physiology, such as heart development,^17^ insulin secretion,^18^ multidrug resistance in acute myeloid leukemia,^19^ endothelial cell adhesion,^20^ cardiac lineage commitment^21^ and epithelial proliferation.^22^ LPHNs are associated with developmental and neurobiological defects,^15^ attention deficit hyperactivity disorder (ADHD)^23^ and cancers.^23–26^ However, the molecular mechanisms underlying LPHN function in development remain unknown.

Latrophilins have several known ligands. In vertebrate neurons, the extracellular domains of LPHNs interact with Teneurins (TENs) in a trans-synaptic manner and regulate excitatory synapse formation and organization.^6, 16, 27–36^ TENs are large cell-surface proteins essential for gastrulation and epithelial to mesenchymal transition (in mice), heart and brain development (in flies), and the correct formation of basement membranes (in worms).^37, 38^ Although, LPHNs and TENs are both key molecules for embryonic development, genetic interaction data for *lat-1* and *ten-1* mutant animals suggested that they act in parallel pathways during embryonic development.^13^ LAT-1 and TEN-1 also appear to be expressed in the same cells, conflicting with a model where interactions between neighboring cells dictate tissue polarity and cell migrations in development. These results altogether implied that LPHNs and TENs may act in parallel on the same cell (*cis*), rather than on neighboring cells (*trans*) and suggested that other yet-to-be-described ligands of LPHNs mediate intercellular interactions for their functions in development.^2^ Vertebrate LPHNs also interact with Fibronectin Leucine Rich Repeat Transmembrane Proteins (FLRTs) to mediate neuronal functions,^39, 40^ but FLRTs do not exist in invertebrates, and thus are likely not involved in the fundamental developmental roles of LPHNs. A recent work reported that *Drosophila* LPHN (Cirl) cooperates with Toll-8 to control planar polarity in embryos.^41^ The various roles of LPHNs in development and the numerous adhesion domains in their extracellular regions further suggest interaction with other ligands.

Here, we aimed to understand the mechanisms of action for LPHNs in their essential but relatively unstudied roles, especially in embryogenesis. To identify proteins that interact with LPHNs, we turned to *C. elegans*, which provides a powerful genetic model to study early development. Importantly, *C. elegans* animals lacking *lat-1* (LPHN homolog) display strong developmental phenotypes. Using our extracellular protein interaction discovery platform in *C. elegans*, we identified a cell-surface receptor, TOL-1 (Toll-like receptor/TLR), the sole Toll-like receptor in *C. elegans*, as a novel binding partner for LAT-1. *tol-1* has similar embryonic phenotypes as *lat-1*. To test whether Toll-like receptors mediate LPHN functions such as embryogenesis, in animals, we employed a multidisciplinary approach to study the function of this interaction: We determined the high-resolution crystal structure of the LAT-1–TOL-1 complex that is consistent with the cryo-EM density map of the full-ectodomain complex, and engineered point mutants at the binding interface to break the complex *in vitro*. We then used CRISPR/Cas9 and generated worms that carry homozygous alleles of *tol-1* or *lat-1* with point mutations that specifically disrupt the LAT-1–TOL-1 interaction. These genetic manipulations in *C. elegans* resulted in developmental defects similar to the ones observed in *tol-1* and *lat-1* null mutant animals. Our study suggests that the TOL-1–LAT-1 interaction is critical for early development in *C. elegans,* and likely in all animals.

## RESULTS

### LAT-1 interacts with the Toll-like receptor in *C. elegans*

To reveal novel interactors for LPHNs and study them in a well-established genetic model system, we chose to work with the nematode *C. elegans*, which provides several advantages: (1) Most conserved cell surface receptor families have fewer members in nematodes, allowing for less extensive screens to cover the putative interaction space. (2) There are only two *C. elegans* LPHN genes, *lat-1* and *lat-2*, but only *lat-1* mutants show major developmental phenotypes, circumventing potential redundancy issues as *lat-2* appears unable to compensate for genetic loss of *lat-1*. (3) There is strong functional conservation to vertebrate LPHNs, where known phenotypes in nematodes match those in mice, including embryonic lethality. (4) Finally, embryonic development is easily tractable in *C. elegans*, compared to mammals.

We included the LAT-1 ectodomain in a protein interaction screen designed to discover extracellular interactions. The high-throughput assay, the Extracellular Interactome Assay (ECIA), uses an ELISA-like setup, where Fc-tagged bait are captured on solid support, and Alkaline phosphatase (AP)-tagged prey report on binding to the bait using chromogenic AP substrates (**Supplementary Fig. 1A**).^42–44^ Crucial to the sensitivity of the assay are high-density display of the bait on plastic support, matched by the oligomerization of the prey using a pentameric coiled coil, resulting in avidity and a large increase in sensitivity of the assay.^43, 45^ We tested LAT-1 against a library of 379 *C. elegans* ectodomains including protein domains known to be conserved and important in developmental functions across bilaterians (see supporting paper available upon request). The LAT-1-Fc bait captured TOL-1-AP prey with the highest significance across the entire library (**Fig. 1B**). Similarly, when TOL-1-Fc was used as bait, LAT-1-AP gave the strongest binding signal (**Fig. 1C**). TOL-1 is a cell surface receptor with two extracellular Leucine-Rich Repeat (LRR) domains, one TM helix and a cytoplasmic Toll-Interleukin receptor-1 receptor (TIR) domain (**Fig. 1D**).^46^ Overall, these results identified the Toll-like receptor TOL-1 ECD as a binding partner for LAT-1 ECD.

Existing literature on nematode *lat*-*1* and *tol-1* demonstrate that these two genes may have roles in closely related developmental functions. In *C. elegans*, *tol-1* mutants show phenotypes in early development similar to those of *lat-1* animals.^47^ Null *tol-1(nr2013)* mutant animals have high rates of arrest during embryonic development, high mortality, and infertility. Expression patterns of *tol-1* in embryo suggest early roles in morphogenesis, while expression in adults is mostly neuronal,^47^ similar to *lat-1* in adults.^12^

### The LAT-1–TOL-1 interaction is direct and stoichiometric

To validate the direct interaction between LAT-1 and TOL-1 with orthogonal protein binding experiments, we created baculoviruses that express constructs covering LAT-1 and TOL-1 ECDs and purified these proteins. The full-length ECD of LAT-1, including the Lectin, HormR and GAIN domains, formed a highly stable complex with the full-length ECD of TOL-1 including the first and second LRR domains, as we observed a shift to the left in the elution volume of a mixed TOL-1–LAT-1 sample compared to the purified TOL-1 ectodomain and the LAT-1 ectodomain individually as analyzed by size-exclusion chromatography (SEC) (**Fig. 1E**). The co-elution of both proteins was confirmed by SDS PAGE analysis (**Fig. 1E**). These results confirmed the high-affinity interaction of LAT-1 with TOL-1.

To determine the exact molecular weight of the complex species that is observed on the size exclusion chromatogram, we used multi-angle light scattering (MALS) and calculated the observed molecular weight as 172 kDa (expected 171.1 kDa) corresponding to a 1:1 stoichiometry (**Fig. 1F and Supplementary Fig. 1B**). The data suggests that the complex is composed of one TOL-1 and one LAT-1 molecule, although we cannot exclude the possibility of lower affinity complexes that might not be observed on a size-exclusion run.

### The lectin domain of LAT-1 binds to the C-LRR domain of TOL-1

To identify TOL-1- and LAT-1-binding domains, we generated a series of domain truncations of LAT-1 and TOL-1, respectively (**Fig. 2A**). Using ECIA, we found that LAT-1 fragments comprising the whole ECD, the ECD without the GAIN domain, or the ECD without the hormone-binding and GAIN domains robustly bound to TOL-1, whereas the LAT-1 fragment without the lectin domain (comprising only the hormone-binding and GAIN domains) did not bind to TOL-1 (**Fig. 2B**). Similarly, TOL-1 fragments comprising the whole ECD, or the ECD without the N-terminal LRR-domain robustly bound to LAT-1, whereas the TOL-1 fragment without the C-terminal LRR-domain (comprising only the N-terminal LRR domain) did not bind (**Fig. 2B**). Western blots of the deletion constructs show that there was no significant change in the level of protein expression and secretion from the cells as compared to the wild-type protein (**Supplementary Figs. 2A, B**). Thus, our results indicate that the lectin domain of LAT-1 and the C-terminal LRR domain of TOL-1 are required for the interaction of LAT-1 and TOL-1.

**Figure 2.**
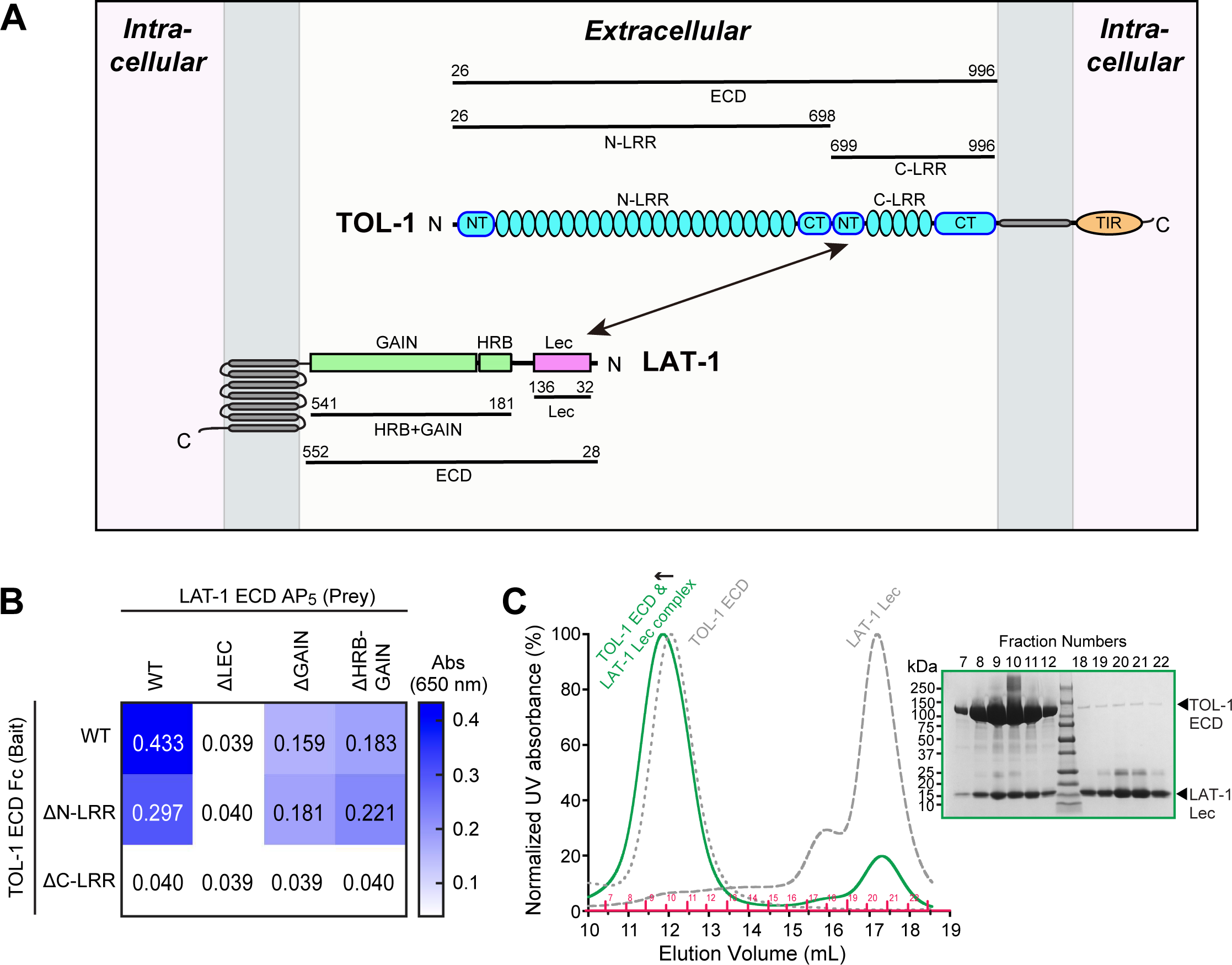
The Lectin domain of LAT-1 and the C-terminal LRR of TOL-1 mediate the LAT-1 –TOL-1 interaction. (A) Schematic of LAT-1 and TOL-1 domain structures and the domain deletion constructs used to map the domains involved in binding. (B) Mapping interacting domains in the LAT-1–TOL-1 complex using ECIA. TOL-1-Fc bait were used to capture LAT-1-AP prey. Binding was detected by absorbance at 650 nm, corresponding to the absorbance of the AP product. Also see Supplementary Fig. 2. (C) The LAT-1 Lectin domain forms a stable complex with the full-length ECD of TOL-1 as shown by SEC profiles and SDS-PAGE analysis of the complex fractions.

To determine whether the lectin domain of LAT-1 and the C-terminal LRR domain of TOL-1 are sufficient for the interaction of LAT-1 and TOL-1, we made baculoviruses that express only the lectin domain of LAT-1 and the C-terminal LRR domain of TOL-1 and purified them separately as recombinant proteins. The lectin domain of LAT-1 formed a stable complex with the full ECD of TOL-1 as analyzed by SEC and SDS-PAGE (**Fig. 2C**). The lectin domain also formed a stable complex with the C-terminal LRR domain of TOL-1 (**Supplementary Fig. 2C**) suggesting that these domains are sufficient for the interaction of LAT-1 and TOL-1. Previous work in *C. elegan*s showed that the Lectin domain is required for rescuing both the lethality and fertility phenotypes of *lat-1* animals.^12^ Our observation that the Lectin domain is the site of TOL-1 binding is consistent with the hypothesis that the tissue polarity and morphogenesis functions of LPHNs are mediated by interactions with TLRs.

### Crystal structure of the LAT-1–TOL-1 complex reveals an unusual mode of interaction

We next set out to determine the high-resolution structure of TOL-1 in complex with LAT-1. We crystallized the full-extracellular region (ECR) of TOL-1 containing both N- and C-terminal LRR domains (residues D26-S996) in a complex with the lectin domain of LAT-1 (residues T32-P136 in isoform a). We determined the x-ray structure of the complex at 4.0 Å resolution using molecular replacement (**Supplementary Table 1**).

The crystal structure of the TOL-1/Lectin complex is a heterodimer with dimensions of ∼125 Å x 45 Å x 96 Å. TOL-1 receptor is comprised of a large semicircular N-terminal LRR domain with 23 LRR repeats followed by a second but smaller C-terminal LRR domain including 5 LRR repeats (**Fig. 3A, B, and Supplementary Fig. 3A, B**), similar to the structure of *Drosophila* Toll.^48^ Each LRR domain is flanked by an N-terminal cap and a C-terminal cap (**Fig. 3A, B**). The C-terminal LRR domain contains the LAT-1-binding site. The lectin domain of LAT-1 binds to the N-terminal cap of the C-terminal LRR domain of TOL-1 and also makes several contacts with the LRR repeats within the C-terminal LRR domain. Surprisingly, the interaction of the Lectin domain with the convex surface of the TOL-1 C-LRR domain is different from other TLR-ligand complexes, where the ligand binding site is located in the concave surface of the LRR region of most TLRs, including the DmToll and Toll5A receptors.^48, 49^ (**Fig. 3A, B**). The direct interaction of the TOL-1 C-terminal LRR with LAT-1 Lectin domain is consistent with our binding data and with previous studies where the lectin domain was identified as critical for the function of LAT-1 in embryo development (**Fig. 2**).^12^ Conservation analysis of the TOL-1–LAT-1 structure suggests that the interface is conserved (**Supplementary Fig. 3 C, D).** The asymmetric unit of our crystals contain four copies of the complex. The N-terminal regions of the N-terminal LRR in TOL-1 has slightly different conformations in each copy as a result of swivel-like motions, suggesting that the N-terminus of TOL-1 has intrinsic flexibility in solution (**Fig 3C**). We observed eight asparagine residues with N-linked glycans attached in the TOL-1 ectodomain, confirming five of the six predicted N-lined glycosylation sites in the N-terminal LRR domain, and three of the six in the C-terminal LRR.

**Figure 3.**
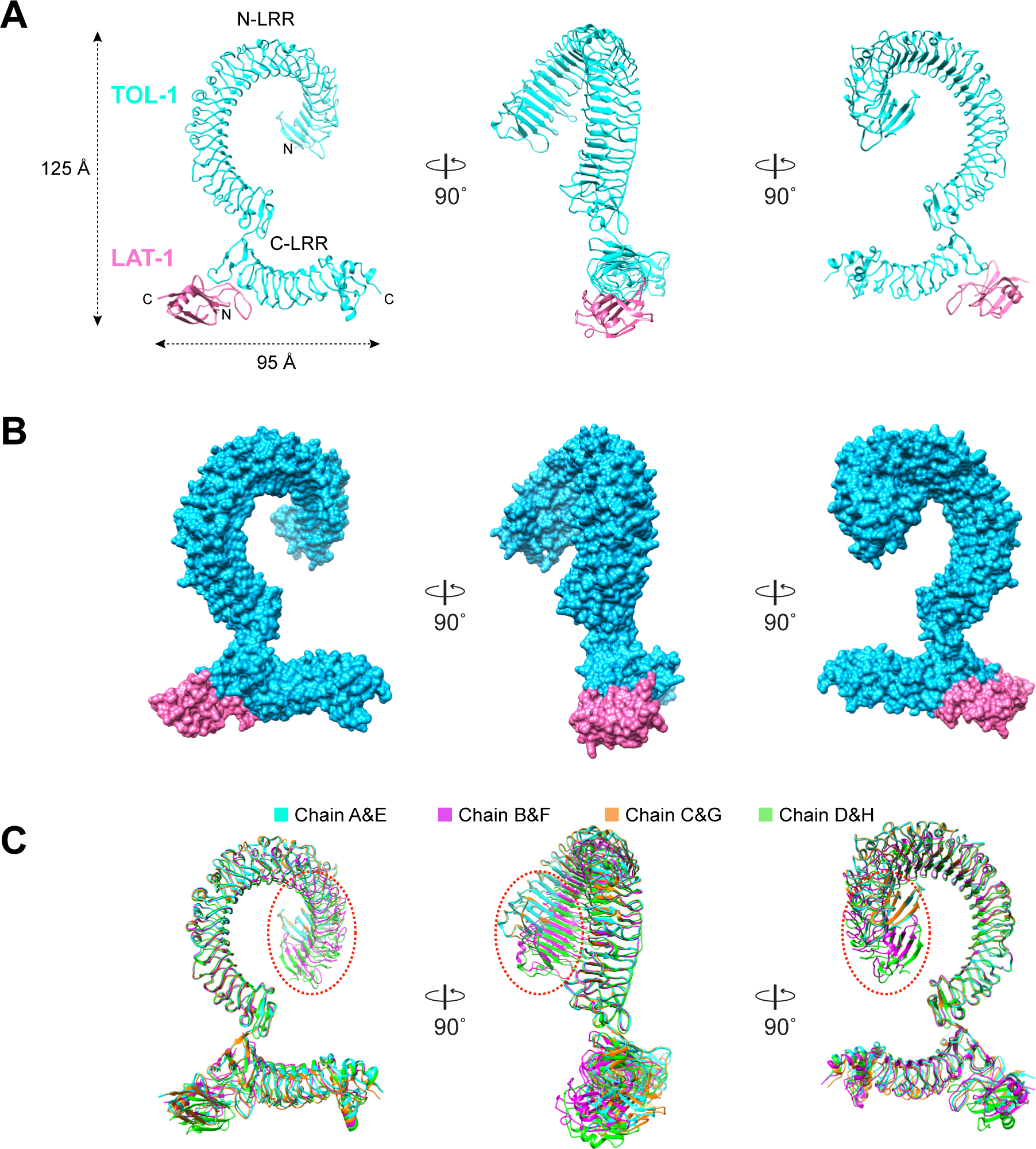
Crystal structure of the LAT-1–TOL-1 complex reveals a previously unobserved mode of interaction (A, B) Crystal structure of the complex of TOL-1 ECD (cyan) with the LAT-1 Lectin domain (pink) (A, cartoon view; B, surface view). (C) Superimposition of the four symmetry elements in the asymmetric unit showing the flexibility of the N-terminal tip of TOL-1.

The LAT-1 lectin domain is similar to previous lectin domain structures from other aGPCRs.^30, 31, 50^ The LAT-1 Lectin domain adopts a kidney shaped structure with overall dimensions of 44 Å x 24 Å x 20 Å, and is composed of five β-strands and a single α-helix. A 24-residue region coming out of the lectin β-sandwich mediates the interaction with TOL-1 C-LRR (**Fig. 3A, B**). We observed no electron density for glycosylation in the lectin domain, as predicted by sequence.

### Cryo-EM density of the LAT-1–TOL-1 complex agrees with the crystal structure

To verify this new mode of interaction, we also collected single-particle cryo-electron microscopy (EM) images of the full ECD TOL-1 in complex with full ECD LAT-1 and obtained a 6.3 Å density map of the full-length complex (**Fig. 4, Supplementary Fig. 4**). 2D class averages showed a similar orientation of LAT-1 with respect to TOL-1 as observed in the crystal structure (**Fig. 4A**). Both the 2D class averages and 3D electron density maps revealed extra density at the N-terminal tip of the second LRR domain (**Fig. 4**), corresponding to the lectin domain of LAT-1. Fitting the crystal structure of the TOL-1–LAT-1 complex into the cryo-EM density map showed excellent correlation and confirmed the validity of the crystal structure (**Fig. 4B**). No density was observed for the HormR and GAIN domains of the LAT-1 molecule likely because there is a 45-amino acid linker between the Lectin and the HormR domains which is expected to introduce flexibility to the molecule. Altogether, our crystallographic structure and cryo-EM maps of the TOL-1–LAT-1 complex are in full agreement.

**Figure 4.**
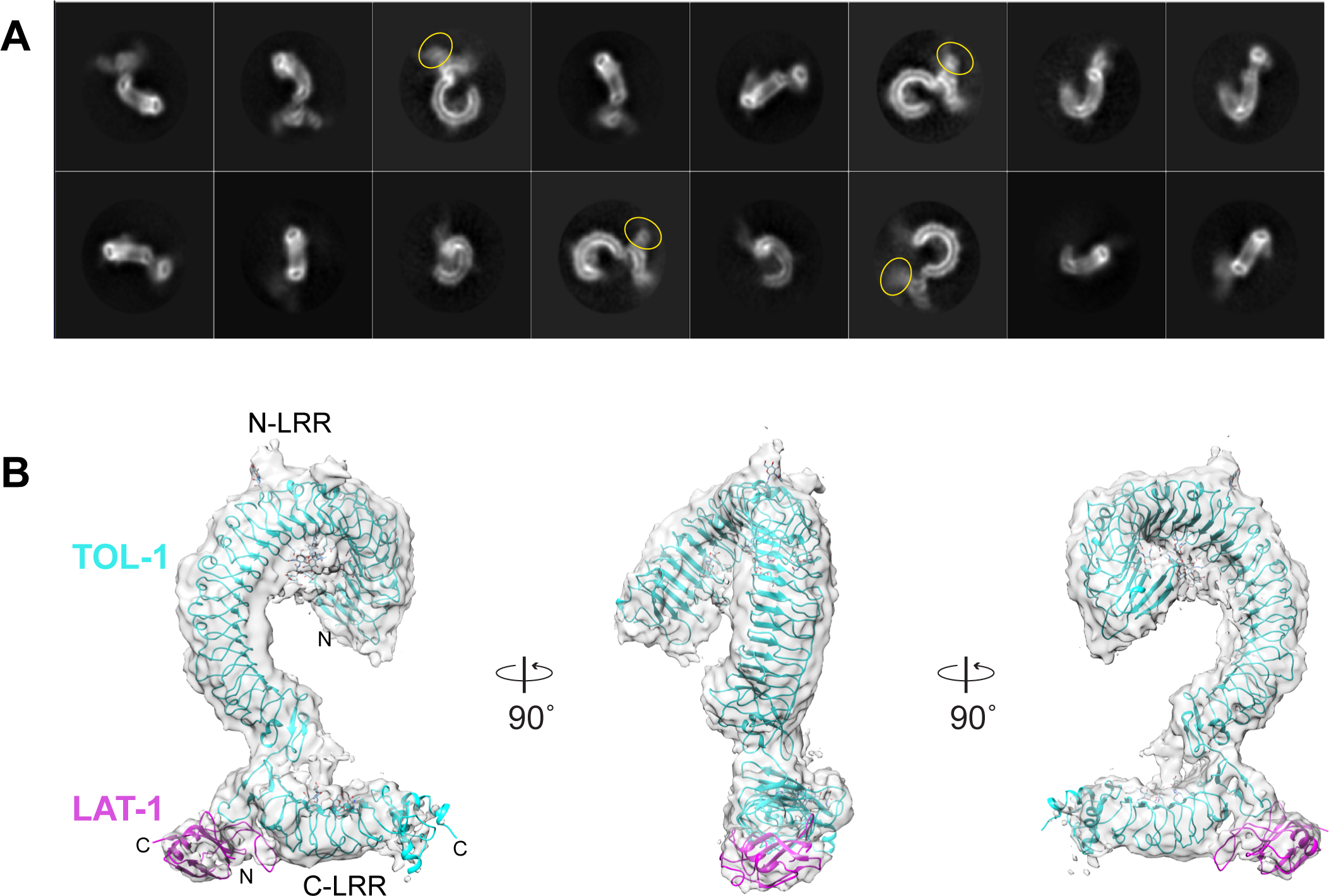
Single-particle cryo-EM density of the full-ECD LAT-1–TOL-1 complex at 6.3 Å resolution. (A) 2-D cryo-EM averages of the TOL-1 ECD–LAT-1 ECD complex. The additional density corresponding to the Lectin domain of LAT-1 is indicated by a yellow circle. The rest of LAT-1 is not visible in averages. (B) 3-D map reconstruction of the TOL-1 ECD–LAT-1 ECD complex with the higher-resolution crystal structure of the complex fit into the cryo-EM density. Also see Supplementary Fig. 4.

### Structure-guided mutagenesis breaks the TOL-1–LAT-1 interaction

The TOL-1–LAT-1 interaction is mediated by an extensive combination of polar and hydrophobic residues from both proteins, creating a large, buried interface area of 1412 Å^2^ (**Fig. 5A, B**). The interface from the crystal structure is consistent with the corresponding cryo-EM electron density (**Supplementary Fig. 5**). The TOL-1 and LAT-1 binding interface is formed by two major interaction points mediated by several interesting structural features from both proteins (**Fig. 5C**, Interaction points are labeled as patch1 and 2 on the figure). The loops from both TOL-1 and LAT-1 extend towards the secondary structure elements in the interacting molecule and make numerous hydrophobic and electrostatic interactions. In patch 1, the N-terminal cap of the C-terminal LRR domain of TOL-1 has a Cys-rich loop with six cysteines that form three disulfide bridges within the loop. These entangled disulfide bridges provide rigidity and enable extension of the loop towards LAT-1 (**Fig. 5D**). In patch 2, on the LAT-1 side, a long 24-amino acid loop comes out of the β-sandwich of the lectin domain and extends towards TOL-1 (**Fig. 5E**). A Cys-residue within this loop makes a disulfide bond with another Cys residue from the β-sandwich providing rigidity and direction to the long loop. This loop interacts with both the N-terminal cap and the leucine-rich repeats of TOL-1 C-LRR. The D79 at the tip of the long loop makes a salt bridge with the K802 on the third leucine-rich repeat of TOL-1.

**Figure 5.**
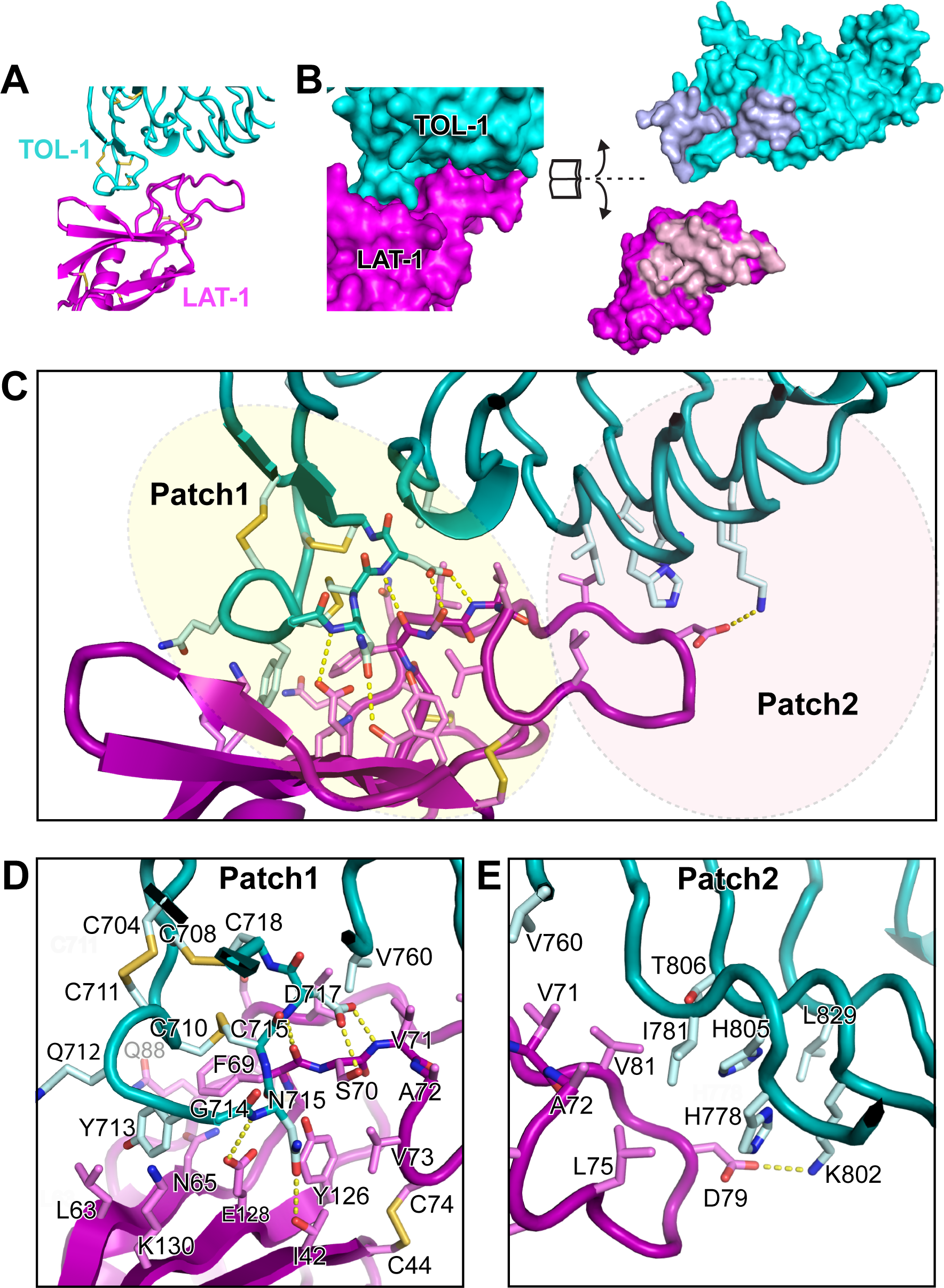
The binding Interface of the LAT-1–TOL-1 complex is large and is mediated by large loops. (A) The structure of the LAT-1–TOL-1 complex binding interface is shown using a cartoon representation, with only the domains involved in the interface used. Disulfide bonds are shown by yellow sticks. (B) (Left) Same as in A but in in surface representation. (Right) Illustrates the buried surfaces using an open book view. Light pink and light blue colors are used to highlight portions of the surfaces involved in the interface. (C) A zoomed in view of the interface in A with side chains shown as sticks. The surface is broadly divided into two patches, each highlighted with a colored oval. (D, E) patches 1, and 2 are shown in detail, with individual residues shown as sticks, with salt bridges and hydrogen bonds shown in yellow dotted lines. See also Supplementary Fig. 5.

To validate the binding interface, we used our crystal structure and sequence conservation analysis to design point mutations on the LAT-1-binding surface of TOL-1 and on the TOL-1-binding surface of LAT-1 that specifically disrupt the interaction. We engineered single- and multiple-point mutations on Fc-tagged TOL-1 and AP_5_-tagged LAT-1, both full ectodomain proteins, and tested the effect of the mutations on the TOL-1–LAT-1 interaction using the ECIA assay. We engineered a TOL-1 ECD mutant (Q712A-Y713A-G714A-N715A) in which we mutated four residues of a LAT-1-binding loop on TOL-1 to Ala (**Fig. 6A**). We also generated several single-site LAT-1 ECD mutants throughout the lectin domain (**Fig. 6A**). Our ECIA assay showed that TOL-1 (Q712A-Y713A-G714A-N715A) mutant has no detectable binding to wild-type LAT-1; and that the LAT-1 F69A mutant has no detectable binding to TOL-1 (**Fig. 6B**). These mutants express and are secreted from S2 cells at levels similar to the wild-type TOL-1 and LAT-1 as confirmed by western blots (Supp Figure). ECIA further confirmed that the second LRR domain of TOL-1 and Lectin domain of LAT-1 are sufficient for binding (**Fig. 6B**). These results show that our cryo-EM and crystallographic models accurately represent the complex structure and provide us with mutants that specifically break their interaction for use in functional assays.

**Figure 6.**
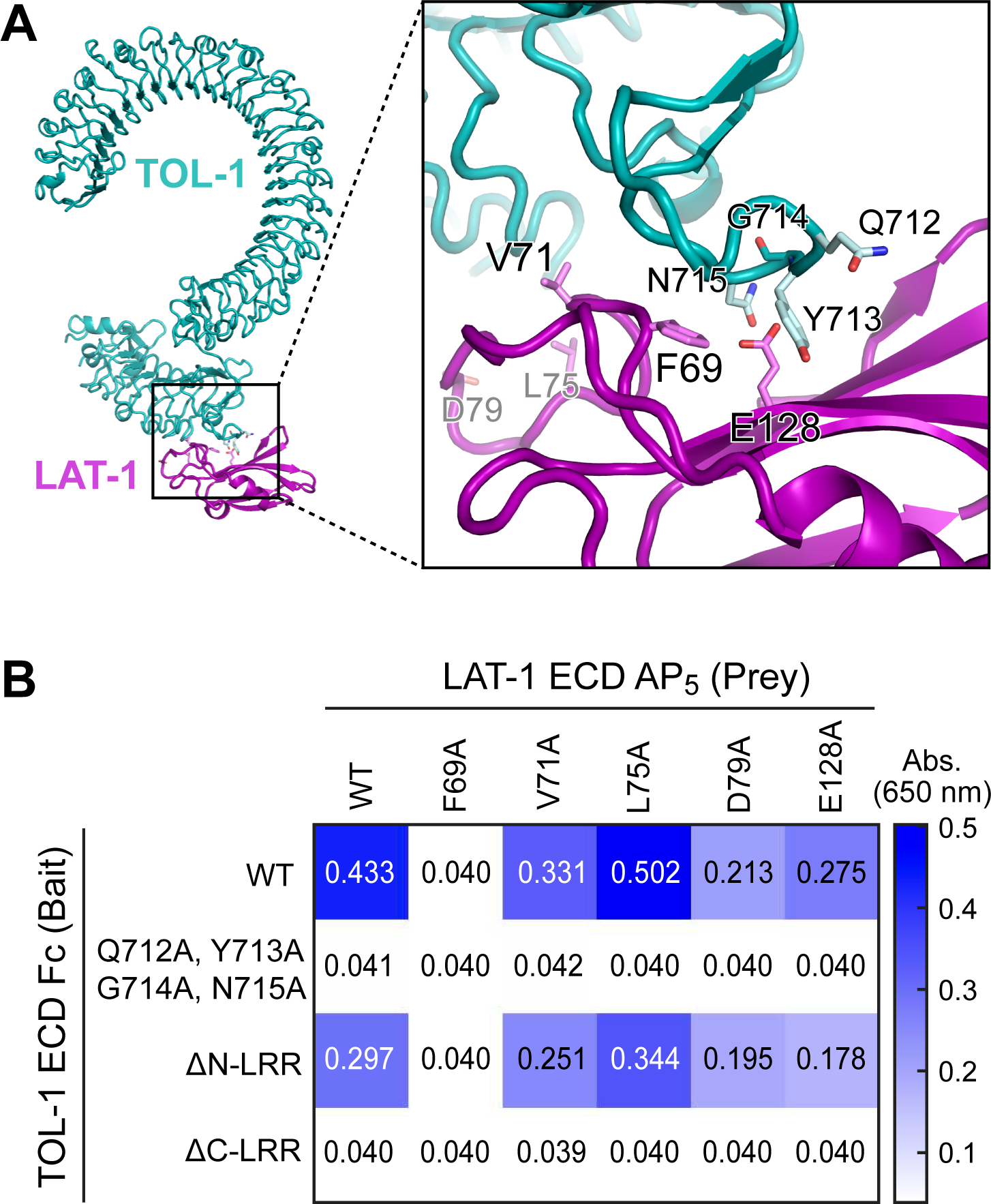
Structure-guided point mutations disrupt the interaction of the LAT-1 and TOL-1. (A) The LAT-1 and TOL-1 interface is shown in a cartoon representation, with key side chains shown as sticks and labeled. (B) ECIA results show that LAT-1 mutations at the TOL-1 interface can diminish or abolish TOL-1 binding. A quadruple TOL-1 mutant designed at the interface also abolishes LAT-1 binding. Also see Supplementary Fig. 2.

### Interaction of TOL-1 and LAT-1 is critical for embryo development and brood size in *C. elegans*

To test whether the LAT-1–TOL-1 interaction is critical for animal development, we undertook a genome editing approach in *C. elegans*. We used CRISPR-Cas9 to generate animals with mutations in *lat-1* and *tol-1* genes designed to selectively break the LAT-1–TOL-1 interaction; we introduced the exact same mutations validated *in vitro* with the ECIA assay (**Fig. 6**). We then tested whether these engineered strains show survival and fertility phenotypes, like the ones described in *lat-1* or *tol-1* strong loss-of-function (putative null) mutant animals. We generated two non-binding mutant strains: the *lat-1(F69A)* strain carries a single point mutation that converts F to A at position 69, whereas the *tol-1(Q712A-Y713A-G714A-N715A)* strain carries four point mutations and is referred to as *tol-1(quad mutant)*. We conducted a phenotypic analysis of these mutants (**Fig. 7A**). Intriguingly, both mutant strains, *lat-1(F69A)* and *tol-1(quad mutant)*, showed defects in brood size and embryo viability (**Fig. 7B**). Like *tol-1* mutant animals carrying a deletion allele (*nr2013*), both the *tol-1(quad mutant)* and the *lat-1(F69A)* animals laid significantly fewer embryos than wildtype (**Fig. 7B**). We also observed embryonic and larval lethality both in the *tol-1(quad mutant)* and *tol-1(nr2013)* animals, which displayed pharyngeal/anterior morphological defects (**Fig. 7C-E**). Consistent with previous work^12^, we observed that *lat-1* mutant animals carrying a deletion allele *(ok1465)* display similar anterior morphological defects (**Fig. 7F**). These results suggest that the direct interaction of TOL-1 and LAT-1 is critical for early development in *C. elegans*.

**Fig. 7.**
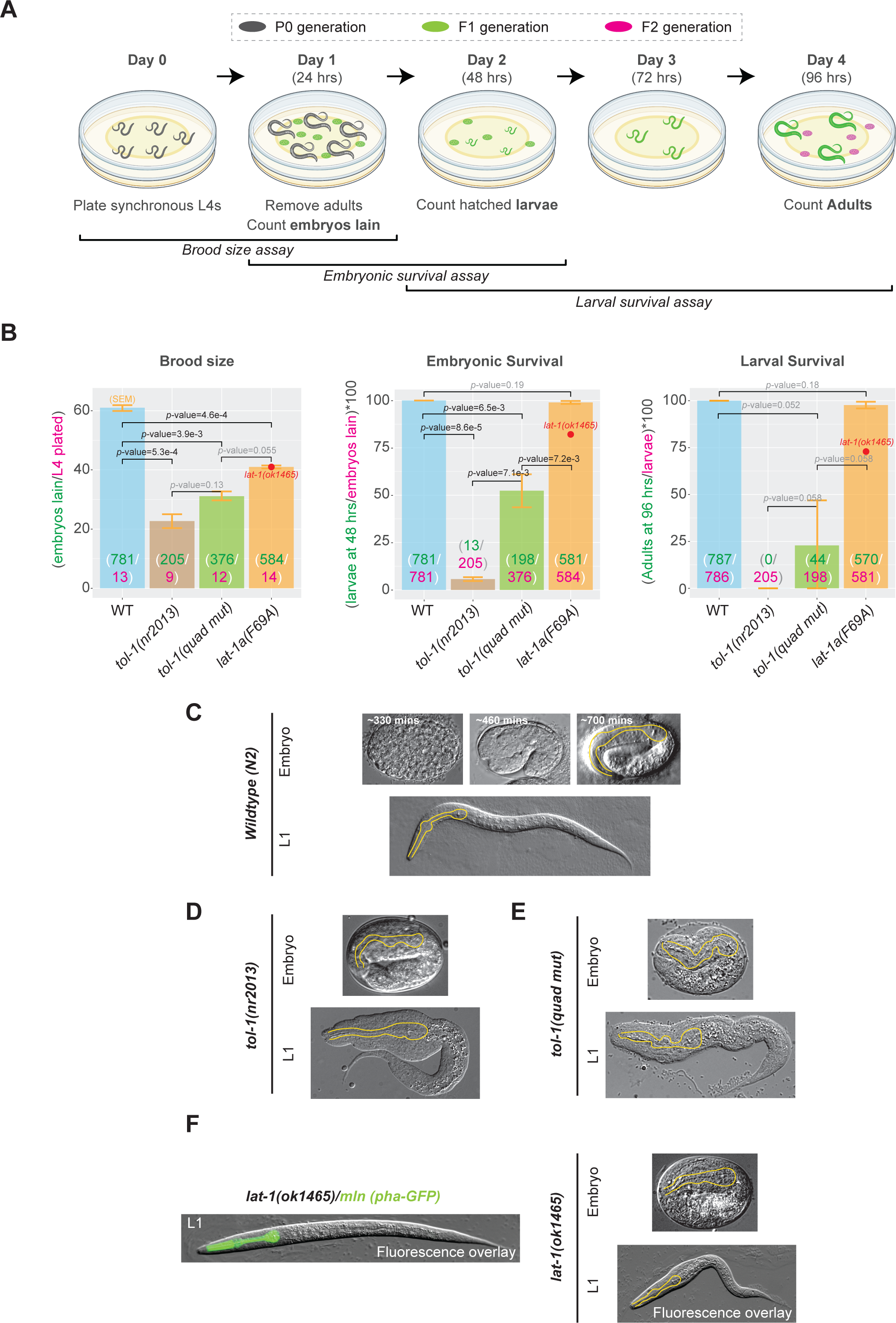
Interaction of TOL-1 and LAT-1 is critical for embryo development and brood size in *C. elegans*. (A) Experimental design for testing the brood size and lethality phenotypes for the engineered *lat-1* and *tol-1* binding mutants. (B) Phenotypes comparing WT, null *tol-1(nr2013)*, *tol-1(quad mut)*, null *lat-1(ok1465)* and *lat-1(F69A)* animals. For brood size (left), data shown are the number of embryos lain per plated L4 animal over the course of 24 hours; For embryonic survival (center), data shown are percentages of embryos that hatch within 24 hours of being lain; for larval survival (right), data shown are percentages of larvae that hatch and survive to adulthood within 94 hours of being lain. Statistical comparisons for brood size, embryonic lethality, and larval lethality of wildtype and each mutant were conducted using a one-tailed t-test (*p*-value < 0.01 are considered significant); exact *p*-values are displayed above each individual comparison; *p*-values in black are significant, whereas *p*-values in grey are not. Error bars refer to the standard error of the mean (SEM). (C) Representative DIC images of wildtype (N2) animals at early, mid, and late stages of embryogenesis (top) and during the first larval stage (L1, bottom). (D-E) Representative DIC images of morphological phenotypes observed in *tol-1(nr2013)* null and *tol-1(quad mutant)* embryos (top) and L1 larvae (bottom). Yellow lines trace the pharyngeal boundaries. (F) Representative DIC-Fluorescence micrograph overlay of heterozygous/balanced *lat-1* L1 larva, labeled with pharyngeal GFP (left) and representative DIC images of morphological phenotypes observed in null *lat-1(ok1465)* embryos (top right) and larvae (bottom right).

## DISCUSSION

While many cell-surface proteins expressed in the developing embryo are known, the receptor-ligand pairings that mediate cellular interactions and regulate developmental functions remain largely enigmatic. Adhesion-GPCRs represent the second largest GPCR family in the human genome, but most adhesion-GPCRs have no known ligands, and the few that have known ligands, such as vertebrate Latrophilins, have more ligands that remain to be identified. While the interaction of vertebrate Latrophilins with TENs and FLRTs are well studied and key for their neuronal functions,^30, 33^ the ligands they interact with to execute essential functions during early embryonic development remained unclear. None of the putative *cis*- or *trans*-interacting ligands for mediating the roles of Latrophilin receptors in embryogenesis were molecularly or structurally characterized as *bona fide* physiological interactions. On the other hand, Toll receptors are a well-studied receptor family with roles in immune system in vertebrates and clear embryonic development defects in invertebrates. Here, we uncovered a convergence between the Latrophilin GPCR and the single Toll-like receptor in *C. elegans*, determined their high-resolution complex structure and revealed a functional receptor-ligand pair in embryonic development. These findings indicate that LAT-1 and TOL-1 bind directly to carry out their critical roles in *C. elegans* development and suggest possible functions for them in higher eukaryotes.

Our data lead to several major conclusions. First, we discovered that LAT-1 adhesion-GPCR forms a direct complex with another signaling receptor, TOL-1 (**Figs. 1, 2, 3, and 4**). TOL-1 is the sole Toll-like receptor in *C. elegans*. Toll-like receptors (TLRs) are critical immunological agents because of their important roles in innate immunity against invading pathogens.^51^ TLR signaling involves a family of adaptor proteins that induce downstream protein kinases which then lead to the activation of transcription factors such as nuclear factor-κB (NF-κB) and the interferon (IFN)-regulatory factor (IRF) family. The key signaling domain, which is unique to the TLR system, is the Toll/interleukin-1 (IL-1) receptor (TIR) domain, which is located in the cytosolic face of each TLR. This domain also exists in the *C. elegans* TOL-1 and is likely the domain used by TOL-1 to initiate intracellular signaling events during embryonic development.^52^

Second, The TOL-1 and LAT-1 interaction is compatible with both a *cis* and *trans* configuration (**Fig. 8**). Positioning the C-termini of the proteins towards the membrane and localizing the complex structure in between two neighboring cell membranes suggests that the proteins can interact in both *cis* and *trans*. Previous studies showed that the vertebrate LPHN3 is a receptor for TEN2 in promoting synapse formation.^33^ Whether the *C. elegans* homolog of TENs, TEN-1, interacts with LAT-1 has not been tested experimentally; and based on genetic studies, this interaction may not exist in *C. elegans*.^2^ Interestingly, the Lectin domain is the binding domain for both TEN-1 and TOL-1. Superimposition of the vertebrate LPHN3–TEN2 complex structure with our invertebrate LAT-1–TOL-1 structure reveals that the Lectin domain uses a different interface for mediating these interactions. Thus, mutagenesis of the TEN-1 binding site on LAT-1 would not interfere with the interaction of LAT-1 with TEN-1 if this interaction were to exist in *C. elegans* (Supplementary Fig. 6). Of note, although Teneurin and TLR binding sites on Latrophilins are distinct and separate, a trimeric complex would not be able to form due to clashes between these two large proteins upon binding to LAT-1. An important question for future studies is whether invertebrate TEN-1 interacts with LAT-1, which could regulate the LAT-1–TOL-1 interaction.

**Fig 8.**
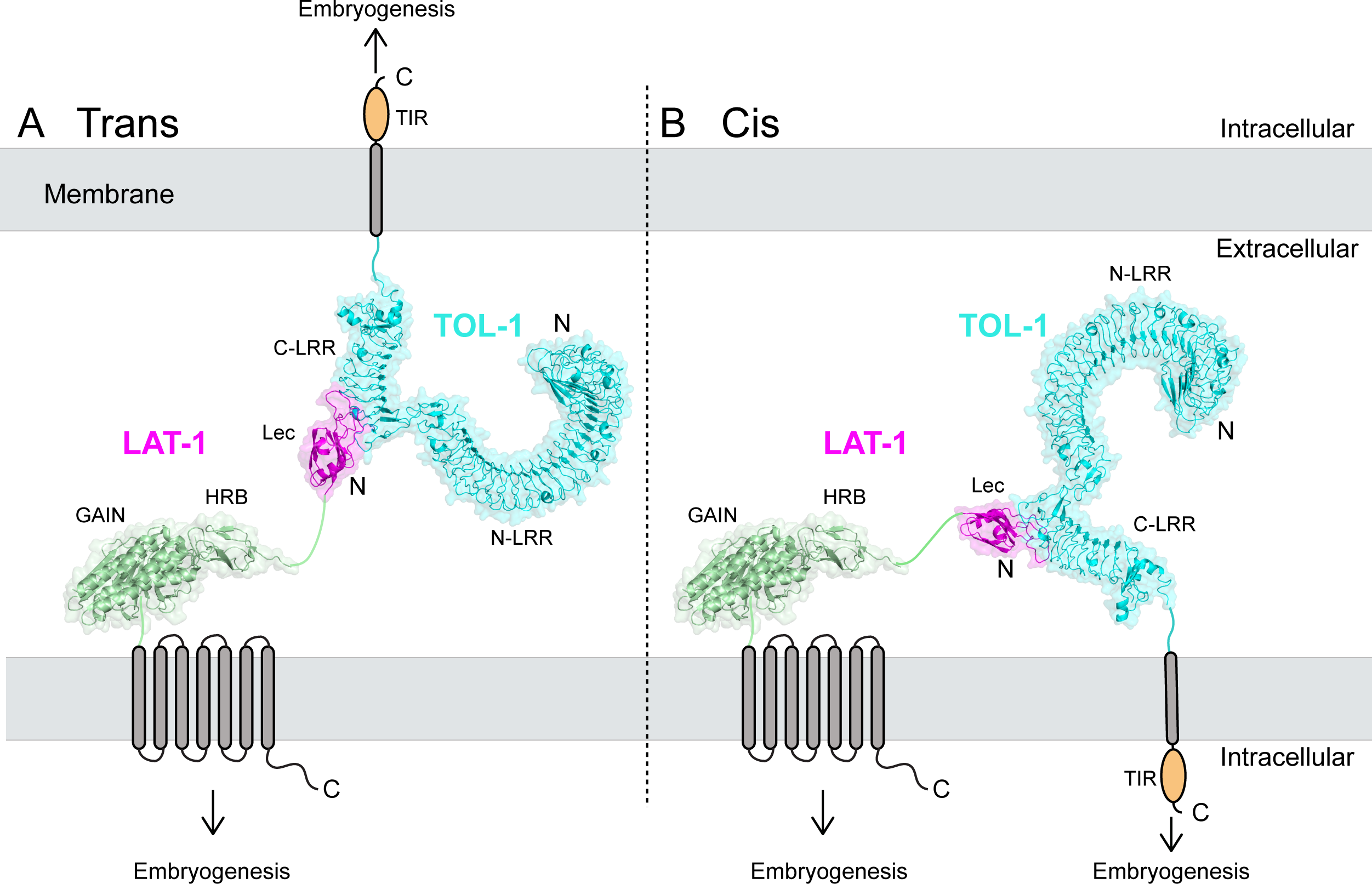
Model showing the interaction of LAT-1 and TOL-1 is important for embryo development The LAT-1–TOL-1 complex identified in this work is compatible with both a *trans* interaction mode, A, and a *cis* interaction mode, B. Each interaction mode may lead to key functions of both LAT-1 and TOL-1 in embryo development. (A) The *trans* interaction mode is the more likely mode for the LAT-1 and TOL-1 interaction. The intracellular TIR domain of TOL-1 and the intracellular G-protein coupling of LAT-1 might be involved in intracellular signal transduction into the developing embryo. Two opposing cell membranes are shown with LAT-1 and TOL-1 molecules projecting out of them, with intracellular faces towards the top and bottom of the page, and an extracellular space in the middle of the page. LAT-1 and TOL-1 are shown as a cartoon representation, linked to their transmembrane helices (shown as rounded cylinders). (B) The *cis* interaction mode is less likely but possible. It might also have a role in regulating the occurrence of the *trans*-interaction to control biological function.

Third, we observed that TOL-1 performs multiple, likely sequential, functions in embryonic development, which are dependent on the TOL-1–LAT-1 interaction. Using CRISPR/Cas9 genome editing in *C. elegans*, the animals that carry point mutations in *tol-1* that abolish the interaction of TOL-1 with LAT-1 phenocopied the *tol-1* null animals, suggesting that the observed phenotypes in *tol-1* null animals are caused by the lack of interaction of TOL-1 with LAT-1 (**Fig. 7A-C**). On the other hand, animals that carry point mutations in *lat-1* that abolish the interaction of LAT-1 with TOL-1 phenocopied only the brood size phenotype of the *lat-1* null animals suggesting that not all phenotypes observed for the *lat-1* null animals are caused by the lack of the LAT-1–TOL-1 interaction (**Fig. 7**). This is most likely due to Latrophilins having multiple binding partners, and demonstrates the power of our structure-based mutagenesis approach to break specific ligand-receptor interactions while maintaining the others *in vivo*.

Finally, our study poses the question of whether Latrophilin-TLR interactions are present in other taxa, especially in flies and vertebrates. In humans, there are three Latrophilins and 10 TLRs, and it is possible that specific sets of human LPHNs and TLRs may interact. It is also possible that the *Drosophila* Latrophilin, Cirl, not only interacts with Toll-8 (Tollo), as it was very recently proposed by Lavalou et al.,^41^ but also with other paralogs among the nine-member fly TLR family. This has the potential to find novel functions for a Latrophilin-TLR interaction in other model organisms. Wormbase reports TOL-1 Is an ortholog of several human genes including CHAD (chondroadherin); ELFN2 (extracellular leucine rich repeat and fibronectin type III domain containing 2); and TRIL (TLR4 interactor with leucine rich repeats). Further studies will reveal whether an orthologous interaction exists in vertebrates.

In summary, our data suggest that the direct interaction of the extracellular regions of LAT-1 and TOL-1 is key for embryo development in *C. elegans*, and its absence leads to partial sterility, lethality, and malformed embryos. LAT-1 and TOL-1 likely exert these functions through *trans*-cellular interactions which may activate downstream signaling pathways of LAT-1 and TOL-1. We present the structure of this complex, which reveals an unexpected molecular architecture, previously not observed in interactions of related binding modules. Since there are three LPHNs, 10 TLRs and numerous LRR repeat containing homologous proteins in mammals, it seems likely that the *trans*-cellular regulatory mechanisms critical for early embryo development observed here may broadly apply to other organisms. This work provides the foundation to reveal new axes of cellular signaling and coordination between two receptor families in the developing embryo, pioneering the identification of homologous pathways in mammals.

## Author Contributions

E.Ö., D.A. and G.C.R. designed all experiments and interpreted results. G.C.R. and J.L. cloned, expressed and purified proteins, carried out bioinformatic and biochemical characterizations and performed x-ray crystallography experiments (with assistance from E.B. and W.I.N.). E.Ö. and G.C.R. collected crystallographic data and performed structure determination, model building and refinement. J.L. performed cryo-EM data collection and map calculation, and refinement (with assistance from M.Z.). P.K. and J.S. designed and performed *C. elegans* experiments. S.C. performed interaction screening experiments (with assistance from E.B.). E.Ö. and D.A. wrote the manuscript with assistance from all authors.

## Data Availability

The coordinates for the crystal structure of LAT-1–TOL-1 crystal structure generated in this study have been deposited in the Protein Data Bank (http://www.rcsb.org) under accession code PDB ID 8SUF. The data for the cryo-EM dataset will be deposited to the EMDB. The authors declare that all data supporting the findings of this study are available within the article and the source data will be provided as a Source Data file.

## Acknowledgements

We thank all members of the Özkan lab for helpful discussions. The work is partly based on research conducted at the Northeastern Collaborative Access Team (NE-CAT) beamlines, which are funded by the National Institute of General Medical Sciences from the National Institutes of Health (NIH) (P30 GM124165). The Eiger 16M detector on the 24-ID-E beam line is funded by an NIH-ORIP HEI grant (S10OD021527). This study used resources of the Advanced Photon Source, a U.S. Department of Energy (DOE) Office of Science User Facility operated for the DOE Office of Science by Argonne National Laboratory under Contract No. DE-AC02-06CH11357. We thank the staff at NE-CAT for excellent support. This work was supported by the grant R35 GM148412 (to D.A.) from NIH.

## Supplementary Figure Legends

**Supplementary Figure 1. Interaction discovery Assay and MALS experiments**

(A) Schematic for the Extracellular Interactome Assay used to discover the LAT-1–TOL-1 interaction (Özkan et al., *Cell*, 2013).

(B) Detailed MALS results for the complex of full ECD of TOL-1 and full ECD of LAT-1.

**Supplementary Figure 2. ECIA assay and western blot of WT and mutant TOL-1 and LAT-1.**

(A) The image of the ECIA plate (Protein A coated plates after bait, prey and AP substrate incubations) after 10 minutes when 650 nm-absorbance is already saturated for WT proteins, showing the mutants have no binding even at saturation.

(B) Comparison of expression of mutant and wild-type TOL-1-Fc and LAT-1-AP proteins as secreted from S2 cells. All bait and prey proteins are C-terminally tagged with hexahistidine tags. Western blot analysis of conditioned media using anti-pentaHis antibodies show comparable expression levels and suggest proper folding of the mutants. LAT-1 constructs including the GAIN domain appear smaller than they are, as the GAIN domain is autoproteolyzed at the GPS cleavage site. This expected modification is known not to result in separation of the cleaved fragments, but causes a shorter C-terminal band to appear on western blots.

(C) The LAT-1 Lectin domain forms a stable complex with the C-terminal LRR domain of TOL-1 as shown by SEC chromatogram and SDS-PAGE analysis of fractions.

**Supplementary Figure 3. Crystallization of TOL-1–LAT-1 complex**

(A) Representative image of the TOL-1–LAT-1 complex crystals used for x-ray data diffraction.

(B) SDS-PAGE gel analysis of the crystals show the presence of both LAT-1 and TOL-1 within the crystals.

(C) The interacting surfaces in the TOL-1–LAT-1 Lectin complex are highly conserved. Coloring reflects conservation between 150 sequences from invertebrates. Also see Supplementary Fig. 3.

(D) Electrostatic potential surface for the TOL-1–LAT-1 complex.

**Supplementary Figure 4. Map Quality and Local Resolution.**

(A) A representative electron micrograph of LAT-1–TOL-1 complex collected using a K3 direct detection camera (Gatan, Inc., Pleasanton, CA). Scale bar is 50 nm.

(B) Fourier shell correlation (FSC) curves of the cryo-EM maps. The resolutions are determined using FSC=0.143 criterion after RELION post-processing. Blue: FSC curve of LAT-1–TOL-1.

(C) Representative 2D class averages of LAT-1–TOL-1 complex. The diameter of the circular mask is 180 Å. The Lec domain is highlighted in yellow circles.

(D) Flow chart of cryo-EM data processing of LAT-1–TOL-1 complex, including particle selection, classification, and 3D classification and refinement. Details are provided in the methods section.

**Supplementary Figure 5. LAT-1–TOL-1 binding interface**

Focused view of the LAT-1–TOL-1 binding interface after fitting the crystal structure of the LAT-1–TOL-1 complex into the single-particle cryo-EM density.

**Supplementary Figure 6.**

Superimposition of the Lectin domain of the LAT-1–TOL-1 complex structure with the Lectin domain of the vertebrate LPHN3–TEN2 complex (PDB ID: 6VHH) reveals that TEN2 and TOL-1 binding sites on the Lectin domain are on different faces of the Lectin domain. However, a trimeric complex formation is not possible because of the clash of TEN with TOL-1 when they are bound. Note that the interaction of *C. elegans* TEN-1 and LAT-1 has not been shown and is believed to not exist.

## METHODS

### Cloning and expression in insect cells

*Caenorhabditis elegans* TOL-1 (Uniprot Q9N5Z3_CAEEL) and LAT-1 (LAT-1, Uniprot LPLT1_CAEEL) constructs were cloned into the pAcGP67A baculovirus transfer vector. Sf9 cells (Thermo Fisher, 12659017) were co-transfected with linearized baculovirus DNA (Expression Systems, 91-002) and the constructed transfer plasmid using the Cellfectin II (Thermo Fisher, 10362100) transfection reagent. Baculoviruses were amplified in Sf9 cells in SF900-III medium containing 10% (v/v) fetal bovine serum (Sigma-Aldrich, F0926). Large-scale protein expression was performed by infection of High Five cells (*Trichoplusia ni*, Thermo Fisher, B85502) in Insect-Xpress medium (Lonza, 12730Q) supplemented with 10 µg/mL gentamicin at a cell density of 2.0 x 10^6^ cells/ml for 72 hours at 27°C.

For structural studies, TOL-1 ECR (residues D26-S996), LAT-1 ECR (residues S28-T552) and LAT-1 Lec domain (residues T32-P136) were cloned with 6xHis-tags. TOL-1 and LAT-1 constructs were co-expressed in High Five insect cells for improved yields. Media containing secreted glycosylated proteins were collected 72 hours after transfection and centrifuged at 900 x *g* for 15 minutes at room temperature. The supernatant was mixed with 50 mM Tris pH 8.0, 5 mM CaCl_2_ and 1 mM NiCl_2_ (final concentrations) and stirred for 30 minutes. After centrifugation at 8,000 g for 30 minutes, the clarified supernatant was incubated with nickel-nitrilotriacetic acid agarose resin (QIAGEN) for 3 hours at room temperature. The resin was washed with HEPES-buffered saline (HBS: 10 mM HEPES, pH 7.2, 150 mM NaCl), supplemented with 20 mM imidazole. The proteins were eluted with HBS containing 200 mM imidazole and further purified using size-exclusion chromatography (Superdex 200 Increase 10/300 or Superose 6 Increase 10/300 columns; GE Healthcare) in 10 mM Tris, pH 8.5, 150 mM NaCl. For the flow cytometry binding assays, LAT-1 Lectin domain (residues T32-P136 in LAT-1a isoform) was cloned with C-terminal 6xHis and Avi tags and purified as described above.

### The extracellular interactome assay (ECIA)

The ectodomains of TOL-1 and LAT-1 were cloned into ECIA expression plasmids both in bait (Fc-tagged) and prey (AP-tagged) forms, and expressed in *Drosophila* S2 cells. The assay was performed as described before.^43^ When engineered mutants of TOL-1 and LAT-1 were tested for binding (Fig. 6B), the expression levels of each mutant series and their wild-type version were normalized for comparison purposes by quantitation via western blots against the 6xHis tags present at the C-termini of all constructs (Antibody: anti-His-Tag Antibody coupled with iFlour 488, Genscript, A01800, 1:500 dilution). Quantitation of fluorescent signal was performed using a Biorad Chemidoc system and ImageLab version 6 (Biorad).

### Multi-angle Light Scattering (MALS)

MALS experiments were done on a Wyatt HELEOS II scattering detector coupled to an ÄKTA FPLC (GE Healthcare) with a Superose 6 Increase 10/300 column using HBS as the sample and running buffer. For data collection and analysis, ASTRA software version 5.3.4 (Wyatt Technology Corp) was used. Average molar mass values calculated using the Zimm model were 169 kDa for the TOL-1–LAT-1 Lec complex (expected heterodimer: 171 kDa).

### X-ray crystallography

Crystallization was achieved using hanging-drop vapor diffusion with 0.2 M sodium malonate, 0.1 M Bis-Tris propane pH 7.5, 26% PEG 3350, 1.2% myo-inositol at 22°C with 16 mg/ml complex. Crystals were cryoprotected in 0.2 M sodium malonate, 0.1 M Bis-Tris propane, pH 7.5, 26% PEG 3350, 1.2% myo-inositol, 30% glycerol. Highly anisotropic diffraction data was collected at the Advanced Photon Source beamline 24-ID-E, and processed using the *XDS* package.^53^ Scaled, unmerged data were ellipsoidically truncated and anisotropically corrected using the *STARANISO* server^54^, which reported a resolution limit of 6.14 Å in the worst direction and 3.55 Å in the best direction, where *I*/σ(*I*) = 1.2 (Table S1).

Molecular replacement was done using *PHASER*^55^ in the *PHENIX* package^56^ with AlphaFold2 models, separately of the TOL-1 first LRR domain, second LRR domain, and the LAT-1 Lectin domain, as predicted using the ColabFold implementation.^57^ *phenix.refine* was used to refine the model structure,^58^ and *COOT* was used for model building and corrections in real space.^59^ For refinement, we used ellipsoidically truncated and anisotropy-corrected structure factors with a high-resolution limit of 4.0 Å, as created by *STARANISO*. Model building and refinement was guided by *MOLPROBITY* tools within *PHENIX* for checking chemical geometry.^60^ Inspection of maps in *COOT* revealed significant swings in the LRR domain “horseshoe” shape, which led us to re-build and rigid-body refine each domain repeat-by-repeat, revealing significant differences between NCS-related copies.

### Cryo-EM data acquisition

3.5 µl purified TOL-1–LAT-1 full-ectodomain complex (0.28 mg/ml) was applied on glow-discharged holey carbon grids (Quantifoil R1.2/1.3, 200 mesh), and vitrified using a Vitrobot Mark IV (FEI Company). The specimen was visualized using a Titan Krios electron microscope (FEI) operating at 300 kV and equipped with a K3 direct electron detector (Gatan, Inc). Images were recorded with a nominal magnification of 81,000x in super-resolution counting mode, corresponding to a pixel size of 0.5315 Å on the specimen level. To maximize data collection speed while keeping image aberrations minimal, image shift was used as imaging strategy using one data position per hole with four holes targeted in one template with one focus position. In total, 3,333 images with defocus values in the range of −1.0 to −2.5 µm were recorded using a dose rate of 14.6 electrons/Å^2^/s. The total exposure time was set to 4.2 s with frames recorded every 0.105 s, resulting in an accumulated dose of about 60.1 electrons per Å^2^ and a total of 40 frames per movie stack.

### Image processing and 3D reconstruction

Stack images were subjected to beam-induced motion correction using MotionCor.^61^ CTF parameters for each micrograph were determined by CTFFIND4.^62^ In total, 1,492,070 particle projections were selected using automated particle picking and subjected to reference-free two-dimensional classification to discard false-positive particles or particles categorized in poorly defined classes, resulting in 1,346,788 particle projections for further processing. Three-dimensional classifications were initially performed on a binned dataset with a pixel size of 2.126 Å using RELION-3.^63^ An ab initio 3D map generated in RELION was used as initial reference model for the first round of maximum-likelihood-based 3D classification. 3D refinement and post-processing was performed on the best class with clear features. The detailed data processing flow is shown in Supplementary Fig 4. Reported resolutions are based on the gold-standard Fourier shell correlation (FSC) using the 0.143 criterion (Supplementary Fig. 4B).

### *C. elegans* brood size and viability assays

Four to five L4 stage-matched hermaphrodite larvae of each genotype were placed on separate 35 mm plates spotted with OP50 or *E. coli* HT115[DE3]. Hermaphrodites were allowed to lay embryos at 25°C for 24 hours. After the 24-hour period, adult worms were removed from plates and total progeny were counted and divided by the number of L4 hermaphrodites originally plated to arrive at brood size number. Progeny at 24 hours were also scored into 2 categories: embryos or larvae. Embryonic survival is expressed as a percentage and was calculated as the number of hatched larvae at 48 hours divided by the total number of embryos laid at 24 hours. Larval survival is expressed as a percentage and was calculated as the number of adults counted at 96 hours divided by the total number of hatched larvae counted at 48 hours.

### C. elegans microscopy

Larval animals were anesthetized with 100 mM sodium azide. Embryonic or larval stage animals were mounted on a 4% agarose pad on imaging slides. Differential interference contrast (DIC) and fluorescent images were taken on an automated fluorescence microscope (Zeiss, Axio Imager Z2) using a Zeiss Axiocam 503 mono (ZEN Blue software, version 2.3.69.1000) equipped with a 63x oil immersion objective. Representative images shown are single z slices. Image reconstruction was performed in Fiji (version 2.9.0/1.53t). Fluorescent overlay and image cropping for Fig. 7 was performed using Adobe Photoshop and Illustrator (23.2.2 Release)

## References

1. Basson, M. A. Signaling in cell differentiation and morphogenesis. Cold Spring Harb. Perspect. Biol. 4, a008151 (2012).

2. Schöneberg, T. & Prömel, S. Latrophilins and Teneurins in Invertebrates: No Love for Each Other? Front. Neurosci. 13, 154 (2019).

3. Topf, U. & Drabikowski, K. Ancient Function of Teneurins in Tissue Organization and Neuronal Guidance in the Nematode Caenorhabditis elegans. Front. Neurosci. 13, 205 (2019).

4. Krasnoperov, V. G. et al. alpha-Latrotoxin stimulates exocytosis by the interaction with a neuronal G-protein-coupled receptor. Neuron 18, 925–937 (1997).

5. Lelianova, V. G. et al. Alpha-latrotoxin receptor, latrophilin, is a novel member of the secretin family of G protein-coupled receptors. J. Biol. Chem. 272, 21504–21508 (1997).

6. Sugita, S., Khvochtev, M. & Südhof, T. C. Neurexins are functional alpha-latrotoxin receptors. Neuron 22, 489–496 (1999).

7. Südhof, T. C. alpha-Latrotoxin and its receptors: neurexins and CIRL/latrophilins. Annu. Rev. Neurosci. 24, 933–962 (2001).

8. Schiöth, H. B., Nordström, K. J. V. & Fredriksson, R. The adhesion GPCRs; gene repertoire, phylogeny and evolution. Adv. Exp. Med. Biol. 706, 1–13 (2010).

9. Hayflick, J. S. A family of heptahelical receptors with adhesion-like domains: a marriage between two super families. J. Recept. Signal Transduct. Res. 20, 119–131 (2000).

10. Araç, D. et al. A novel evolutionarily conserved domain of cell-adhesion GPCRs mediates autoproteolysis. EMBO J. 31, 1364–1378 (2012).

11. Moreno-Salinas, A. L. et al. Latrophilins: A Neuro-Centric View of an Evolutionary Conserved Adhesion G Protein-Coupled Receptor Subfamily. Front. Neurosci. 13, 700 (2019).

12. Langenhan, T. et al. Latrophilin signaling links anterior-posterior tissue polarity and oriented cell divisions in the C. elegans embryo. Dev. Cell 17, 494–504 (2009).

13. Prömel, S. et al. The GPS motif is a molecular switch for bimodal activities of adhesion class G protein-coupled receptors. Cell Rep. 2, 321–331 (2012).

14. Silva, J.-P. & Ushkaryov, Y. A. The latrophilins, ‘split-personality’ receptors. Adv. Exp. Med. Biol. 706, 59–75 (2010).

15. Vitobello, A. et al. ADGRL1 haploinsufficiency causes a variable spectrum of neurodevelopmental disorders in humans and alters synaptic activity and behavior in a mouse model. Am. J. Hum. Genet. 109, 1436–1457 (2022).

16. Anderson, G. R. et al. Postsynaptic adhesion GPCR latrophilin-2 mediates target recognition in entorhinal-hippocampal synapse assembly. J. Cell Biol. 216, 3831–3846 (2017).

17. Lee, C.-S. et al. The G Protein-Coupled Receptor Latrophilin-2, A Marker for Heart Development, Induces Myocardial Repair After Infarction. Stem Cells Transl. Med. 11, 332– 342 (2022).

18. Röthe, J. et al. Involvement of the Adhesion GPCRs Latrophilins in the Regulation of Insulin Release. Cell Rep. 26, 1573–1584.e5 (2019).

19. Kocibalova, Z. et al. Development of Multidrug Resistance in Acute Myeloid Leukemia Is Associated with Alterations of the LPHN1/GAL-9/TIM-3 Signaling Pathway. Cancers 13, 3629 (2021).

20. Camillo, C. et al. LPHN2 inhibits vascular permeability by differential control of endothelial cell adhesion. J. Cell Biol. 220, e202006033 (2021).

21. Lee, C.-S. et al. Adhesion GPCR Latrophilin-2 Specifies Cardiac Lineage Commitment through CDK5, Src, and P38MAPK. Stem Cell Rep. 16, 868–882 (2021).

22. Bucar, E. B., Carmelo, A., Thai, M. H. & Frey, M. R. Loss of Adhesion G-Protein-Coupled Receptor L2 Expression Impacts Colonic Epithelial Proliferation. FASEB J. 36, (2022).

23. Arcos-Burgos, M. et al. A common variant of the latrophilin 3 gene, LPHN3, confers susceptibility to ADHD and predicts effectiveness of stimulant medication. Mol. Psychiatry 15, 1053–1066 (2010).

24. O’Hayre, M. et al. The emerging mutational landscape of G proteins and G-protein-coupled receptors in cancer. Nat. Rev. Cancer 13, 412–424 (2013).

25. Kan, Z. et al. Diverse somatic mutation patterns and pathway alterations in human cancers. Nature 466, 869–873 (2010).

26. Sumbayev, V. V. et al. Expression of functional neuronal receptor latrophilin 1 in human acute myeloid leukaemia cells. Oncotarget 7, 45575–45583 (2016).

27. Meza-Aguilar, D. G. & Boucard, A. A. Latrophilins updated. Biomol. Concepts 5, 457–478 (2014).

28. Silva, J.-P. et al. Latrophilin 1 and its endogenous ligand Lasso/teneurin-2 form a high-affinity transsynaptic receptor pair with signaling capabilities. Proc. Natl. Acad. Sci. U. S. A. 108, 12113–12118 (2011).

29. Sugita, S., Ichtchenko, K., Khvotchev, M. & Südhof, T. C. alpha-Latrotoxin receptor CIRL/latrophilin 1 (CL1) defines an unusual family of ubiquitous G-protein-linked receptors. G-protein coupling not required for triggering exocytosis. J. Biol. Chem. 273, 32715–32724 (1998).

30. Li, J. et al. Alternative splicing controls teneurin-latrophilin interaction and synapse specificity by a shape-shifting mechanism. Nat. Commun. 11, 2140 (2020).

31. del Toro, D. et al. Structural Basis of Teneurin-Latrophilin Interaction in Repulsive Guidance of Migrating Neurons. Cell 180, 323–339.e19 (2020).

32. Nazarko, O. et al. A Comprehensive Mutagenesis Screen of the Adhesion GPCR Latrophilin-1/ADGRL1. iScience 3, 264–278 (2018).

33. Sando, R., Jiang, X. & Südhof, T. C. Latrophilin GPCRs direct synapse specificity by coincident binding of FLRTs and teneurins. Science 363, eaav7969 (2019).

34. Zhang, R. S., Liakath-Ali, K. & Südhof, T. C. Latrophilin-2 and latrophilin-3 are redundantly essential for parallel-fiber synapse function in cerebellum. eLife 9, e54443 (2020).

35. Boucard, A. A., Maxeiner, S. & Südhof, T. C. Latrophilins function as heterophilic cell-adhesion molecules by binding to teneurins: regulation by alternative splicing. J. Biol. Chem. 289, 387–402 (2014).

36. Li, J. et al. Structural Basis for Teneurin Function in Circuit-Wiring: A Toxin Motif at the Synapse. Cell 173, 735–748.e15 (2018).

37. Levine, A. et al. Odd Oz: a novel Drosophila pair rule gene. Cell 77, 587–598 (1994).

38. Trzebiatowska, A., Topf, U., Sauder, U., Drabikowski, K. & Chiquet-Ehrismann, R. Caenorhabditis elegans teneurin, ten-1, is required for gonadal and pharyngeal basement membrane integrity and acts redundantly with integrin ina-1 and dystroglycan dgn-1. Mol. Biol. Cell 19, 3898–3908 (2008).

39. O’Sullivan, M. L. et al. FLRT proteins are endogenous latrophilin ligands and regulate excitatory synapse development. Neuron 73, 903–910 (2012).

40. Lu, Y. C. et al. Structural Basis of Latrophilin-FLRT-UNC5 Interaction in Cell Adhesion. Struct. Lond. Engl. 1993 23, 1678–1691 (2015).

41. Lavalou, J. et al. Formation of polarized contractile interfaces by self-organized Toll-8/Cirl GPCR asymmetry. Dev. Cell 56, 1574–1588.e7 (2021).

42. Bushell, K. M., Söllner, C., Schuster-Boeckler, B., Bateman, A. & Wright, G. J. Large-scale screening for novel low-affinity extracellular protein interactions. Genome Res. 18, 622–630 (2008).

43. Özkan, E. et al. An Extracellular Interactome of Immunoglobulin and LRR Proteins Reveals Receptor-Ligand Networks. Cell 154, 228–239 (2013).

44. Wojtowicz, W. M. et al. A vast repertoire of Dscam binding specificities arises from modular interactions of variable Ig domains. Cell 130, 1134–1145 (2007).

45. Voulgaraki, D. et al. Multivalent recombinant proteins for probing functions of leucocyte surface proteins such as the CD200 receptor. Immunology 115, 337–346 (2005).

46. Botos, I., Segal, D. M. & Davies, D. R. The structural biology of Toll-like receptors. Struct. Lond. Engl. 1993 19, 447–459 (2011).

47. Pujol, N. et al. A reverse genetic analysis of components of the Toll signaling pathway in Caenorhabditis elegans. Curr. Biol. CB 11, 809–821 (2001).

48. Parthier, C. et al. Structure of the Toll-Spätzle complex, a molecular hub in Drosophila development and innate immunity. Proc. Natl. Acad. Sci. U. S. A. 111, 6281–6286 (2014).

49. Saucereau, Y. et al. Structure and dynamics of Toll immunoreceptor activation in the mosquito Aedes aegypti. Nat. Commun. 13, 5110 (2022).

50. Leon, K. et al. Structural basis for adhesion G protein-coupled receptor Gpr126 function. Nat. Commun. 11, 194 (2020).

51. Akira, S., Uematsu, S. & Takeuchi, O. Pathogen recognition and innate immunity. Cell 124, 783–801 (2006).

52. O’Neill, L. A. J. & Bowie, A. G. The family of five: TIR-domain-containing adaptors in Toll-like receptor signalling. Nat. Rev. Immunol. 7, 353–364 (2007).

53. Kabsch, W. XDS. Acta Crystallogr. D Biol. Crystallogr. 66, 125–132 (2010).

54. Tickle, I. J. et al. STARANISO. (2018).

55. McCoy, A. J. et al. *Phaser* crystallographic software. J. Appl. Crystallogr. 40, 658–674 (2007).

56. Liebschner, D. et al. Macromolecular structure determination using X-rays, neutrons and electrons: recent developments in *Phenix*. Acta Crystallogr. Sect. Struct. Biol. 75, 861–877 (2019).

57. Mirdita, M. et al. ColabFold: making protein folding accessible to all. Nat. Methods 19, 679– 682 (2022).

58. Afonine, P. V. et al. Towards automated crystallographic structure refinement with *phenix.refine*. Acta Crystallogr. D Biol. Crystallogr. 68, 352–367 (2012).

59. Emsley, P., Lohkamp, B., Scott, W. G. & Cowtan, K. Features and development of *Coot*. *Acta Crystallogr*. D Biol. Crystallogr. 66, 486–501 (2010).

60. Chen, V. B., et al. *MolProbity*: all-atom structure validation for macromolecular crystallography. Acta Crystallogr. D Biol. Crystallogr. 66, 12–21 (2010).

61. Li, X. et al. Electron counting and beam-induced motion correction enable near-atomic-resolution single-particle cryo-EM. Nat. Methods 10, 584–590 (2013).

62. Mindell, J. A. & Grigorieff, N. Accurate determination of local defocus and specimen tilt in electron microscopy. J. Struct. Biol. 142, 334–347 (2003).

63. Zivanov, J. et al. New tools for automated high-resolution cryo-EM structure determination in RELION-3. eLife 7, e42166 (2018).

