## Supplemental Materials for "Structural basis and functional roles for Toll-like receptor binding to Latrophilin adhesion-GPCR in embryo development"

**Supplementary Material:**

Supplementary Figures 1-6

Supplementary Tables 1

**A**

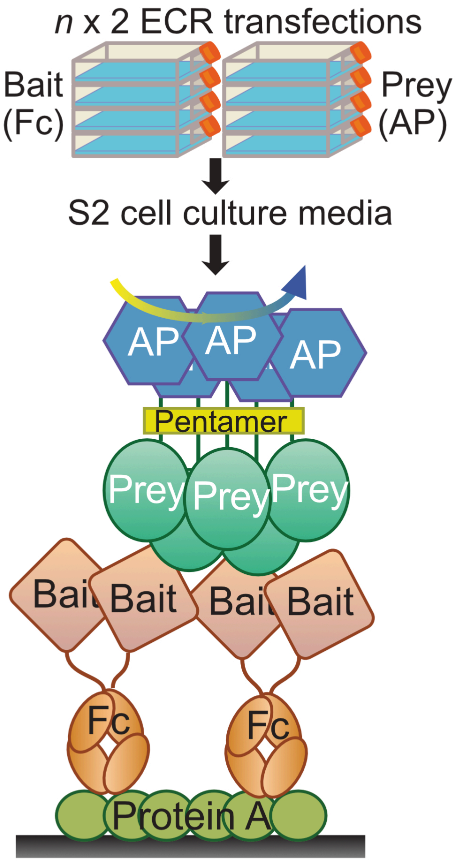


**B**

Peak 1

| | |
|---|---|
| **Polydispersity** | |
| Mw/Mn | 1.000(0.2%) |
| Mz/Mn | 1.000(0.4%) |
| **rms radius moments (nm)** | |
| Rn | 4.6(134%) |
| Rw | 4.6(134%) |
| Rz | 4.6(134%) |
| **Molar mass moments (g/mol)** | |
| Mn | 1.694e+5(0.2%) |
| Mp | 1.696e+5(0.1%) |
| Mw | 1.694e+5(0.2%) |
| Mz | 1.694e+5(0.4%) |

Calculated MW = 172 kDa

**Supplementary Figure 1. Interaction discovery Assay and MALS experiments**
(A) Schematic for the Extracellular Interactome Assay used to discover the LAT-1–TOL-1 interaction (Özkan et al., Cell, 2013).
(B) Detailed MALS results for the complex of full ECD of TOL-1 and full ECD of LAT-1.

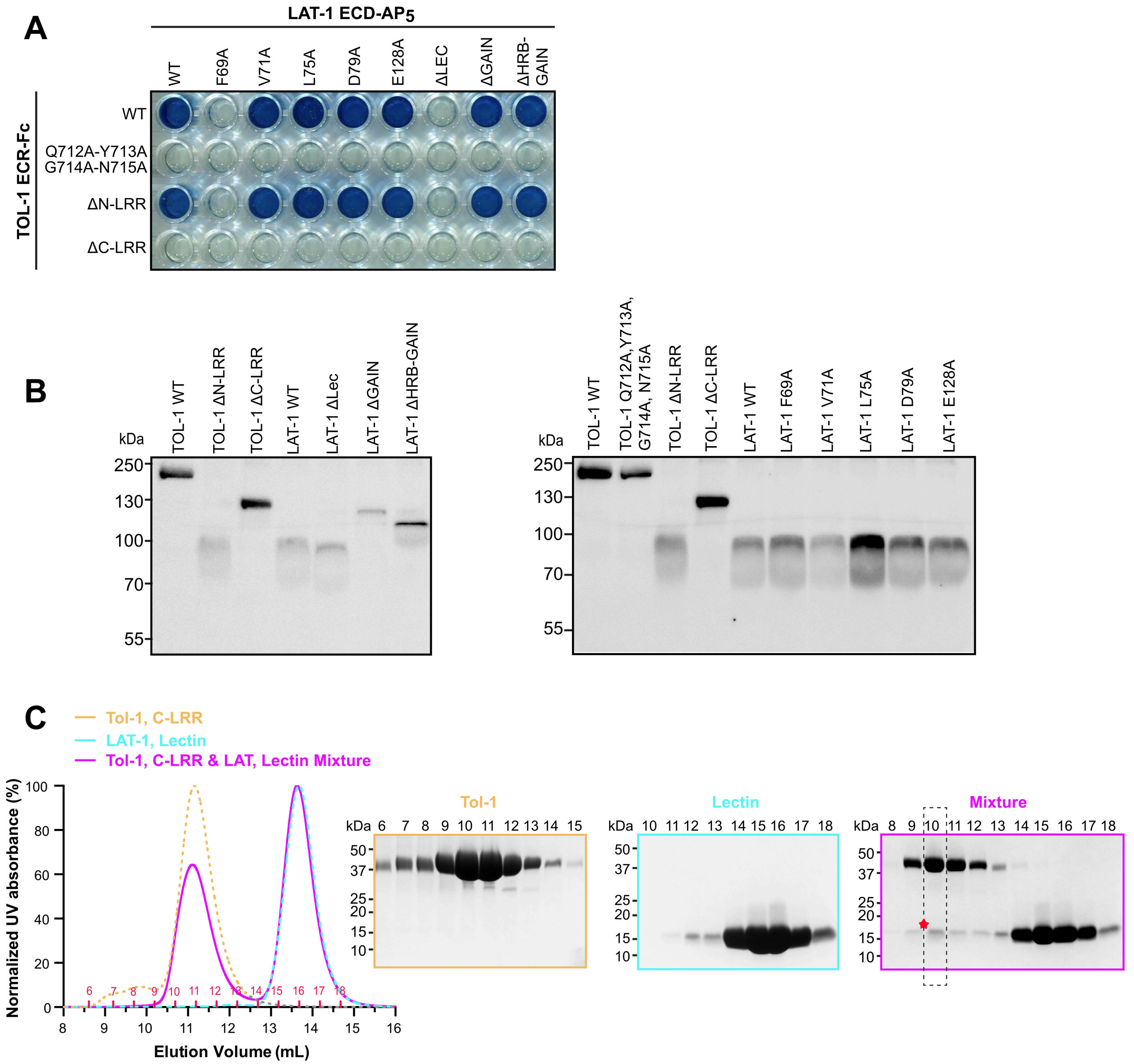


**Supplementary Figure 2. ECIA assay and western blot of WT and mutant TOL-1 and LAT-1.**
(A) The image of the ECIA plate (Protein A coated plates after bait, prey and AP substrate incubations) after 10 minutes when 650 nm-absorbance is already saturated for WT proteins, showing the mutants have no binding even at saturation.
(B) Comparison of expression of mutant and wild-type TOL-1-Fc and LAT-1-AP proteins as secreted from S2 cells. All bait and prey proteins are C-terminally tagged with hexahistidine tags. Western blot analysis of conditioned media using anti-pentaHis antibodies show comparable expression levels and suggest proper folding of the mutants. LAT-1 constructs including the GAIN domain appear smaller than they are, as the GAIN domain is autoproteolyzed at the GPS cleavage site. This expected modification is known not to result in separation of the cleaved fragments, but causes a shorter C-terminal band to appear on western blots.
(C) The LAT-1 Lectin domain forms a stable complex with the C-terminal LRR domain of TOL-1 as shown by SEC chromatogram and SDS-PAGE analysis of fractions.

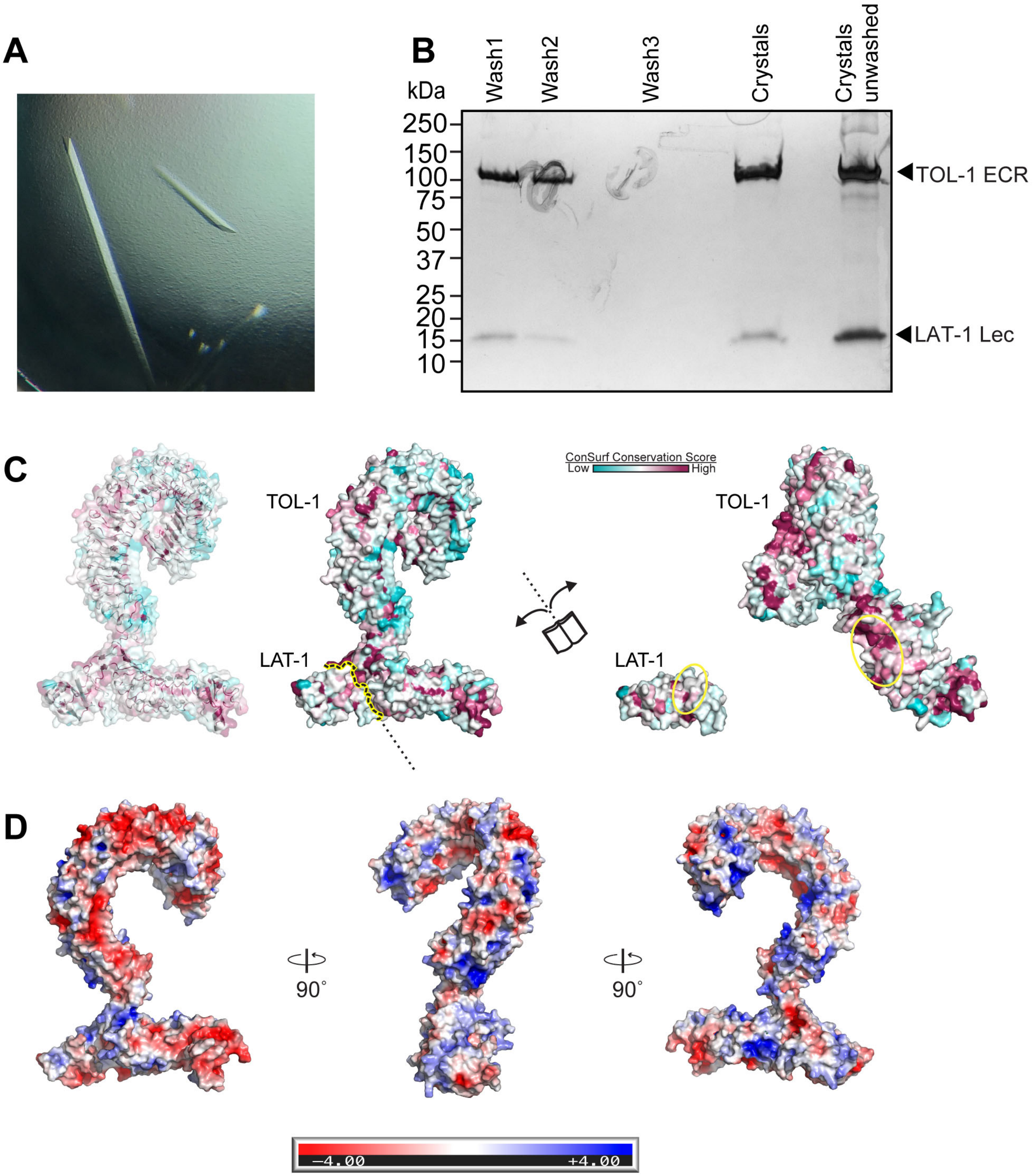


**Supplementary Figure 3. Crystallization of TOL-1–LAT-1 complex**
(A) Representative image of the TOL-1–LAT-1 complex crystals used for x-ray data diffraction.
(B) SDS-PAGE gel analysis of the crystals show the presence of both LAT-1 and TOL-1 within the crystals.
(C) The structure of the TOL-1–LAT-1 complex is shown in surface representation on which the conservation of residues is mapped from most conserved (magenta) to least conserved (cyan) using the ConSurf server. Yellow dotted lines enclose the TOL-1–LAT-1 binding interface. The LAT-1 binding site on TOL-1 and the TOL-1 binding site on LAT-1 are indicated by yellow circles.
(D) Electrostatic potential surface for the TOL-1–LAT-1 complex.

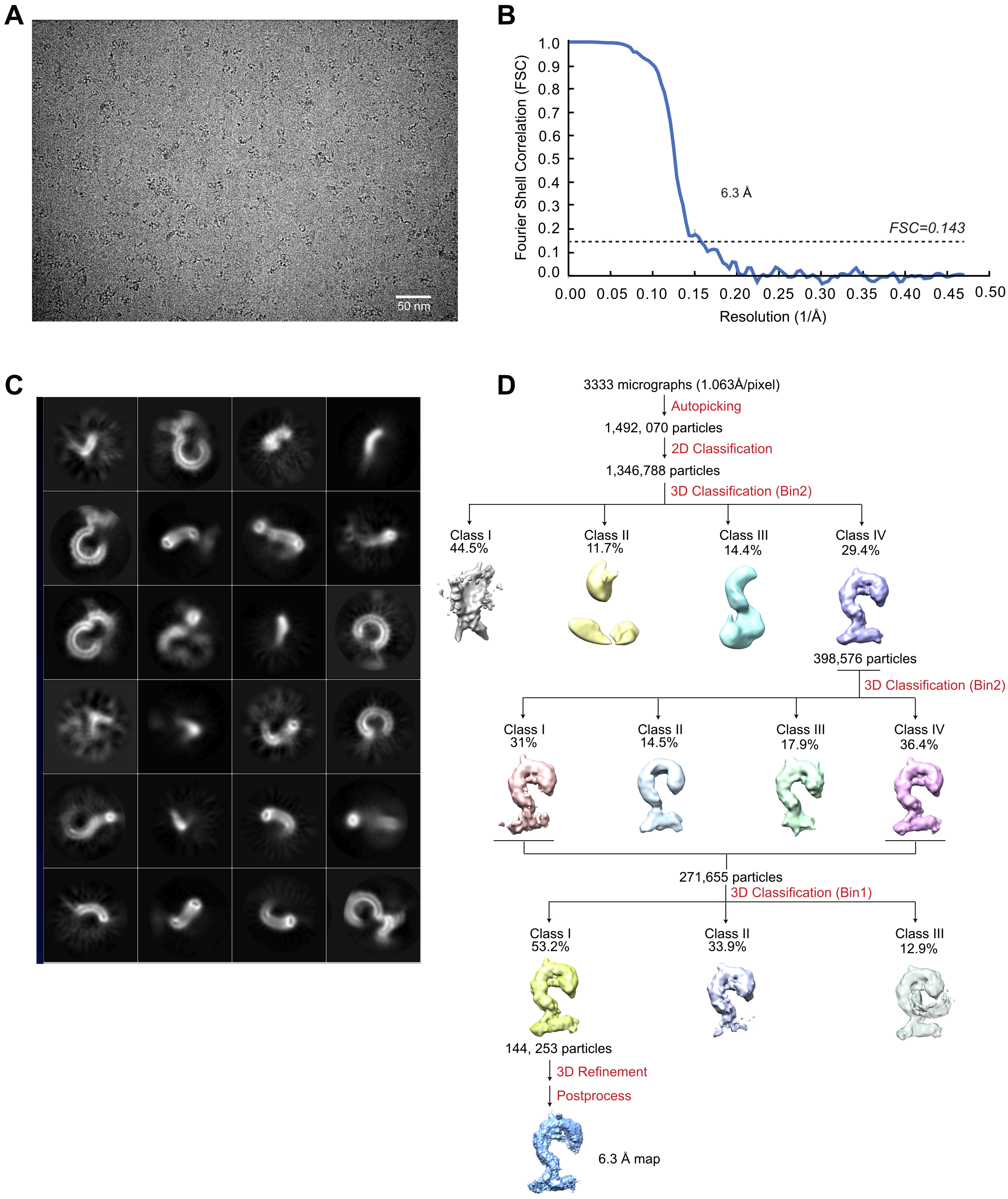


**Supplementary Figure 4. Map Quality and Local Resolution.**

(A) A representative electron micrograph of LAT-1–TOL-1 complex collected using a K3 direct detection camera (Gatan, Inc., Pleasanton, CA). Scale bar is 50 nm.

(B) Fourier shell correlation (FSC) curves of the cryo-EM maps. The resolutions are determined using FSC=0.143 criterion after RELION post-processing. Blue: FSC curve of LAT-1–TOL-1.

(C) Representative 2D class averages of LAT-1–TOL-1 complex. The diameter of the circular mask is 180 Å. The Lec domain is highlighted in yellow circles.

(D) Flow chart of cryo-EM data processing of LAT-1–TOL-1 complex, including particle selection, classification, and 3D classification and refinement. Details are provided in the methods section.

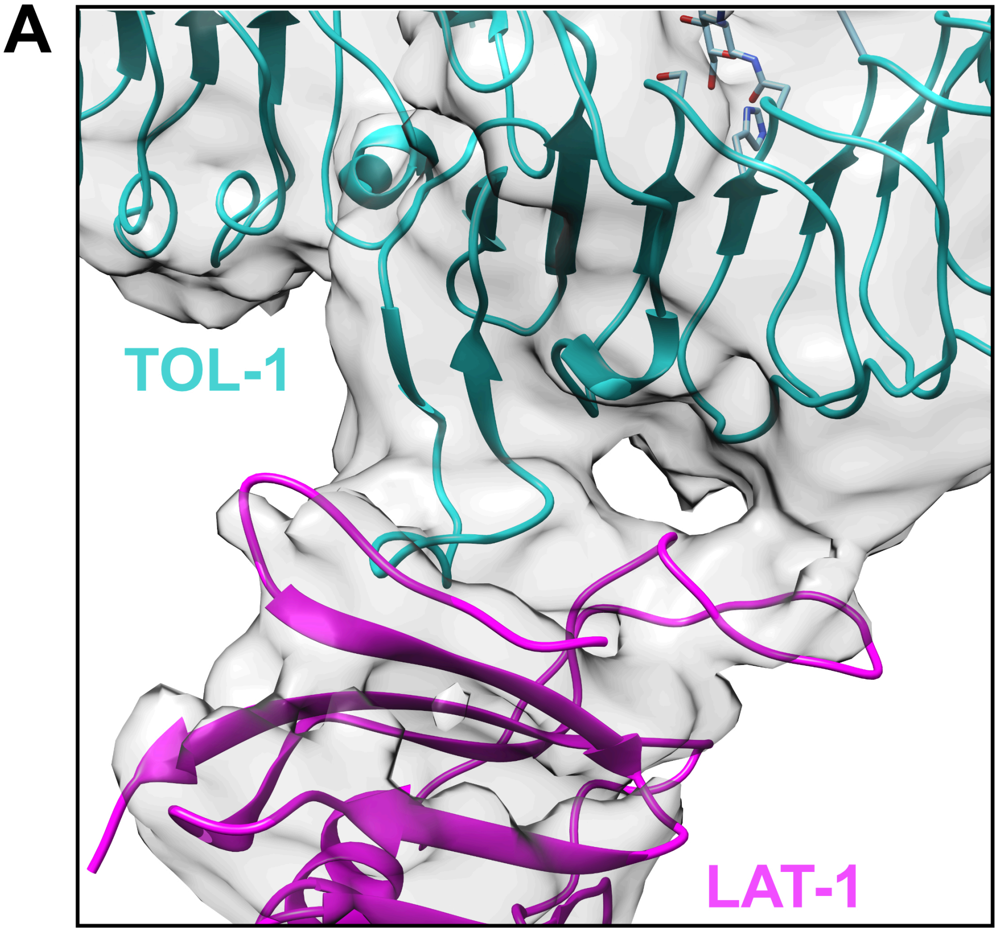


**Supplementary Figure 5. LAT-1–TOL-1 binding interface**
Focused view of the LAT-1–TOL-1 binding interface after fitting the crystal structure of the LAT-1–TOL-1 complex into the single-particle cryo-EM density.

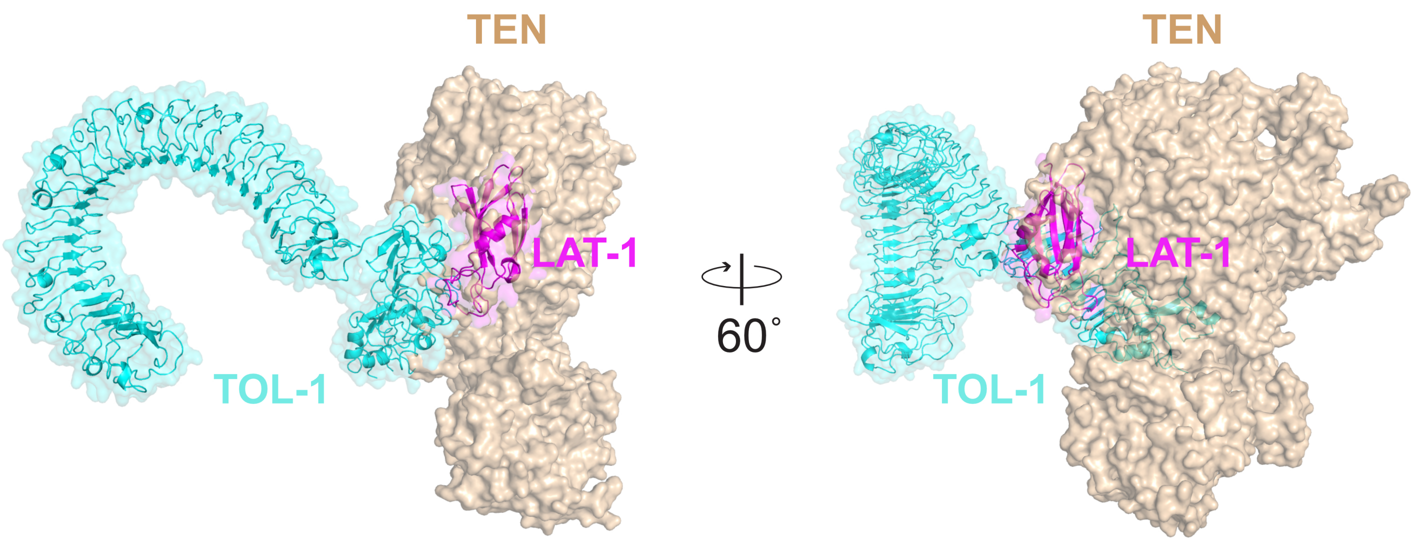


**Supplementary Figure 6.**
Superimposition of the Lectin domain of the LAT-1–TOL-1 complex structure with the Lectin domain of the vertebrate LPHN3–TEN2 complex (PDB ID: 6VHH) reveals that TEN2 and TOL-1 binding sites on the Lectin domain are on different faces of the Lectin domain. However, a trimeric complex formation is not possible because of the clash of TEN with TOL-1 when they are bound. Note that the interaction of *C. elegans* TEN-1 and LAT-1 has not been shown and is believed to not exist.

**Table S1. Data and refinement statistics for x-ray crystallography.**

| | TOL-1 ECD + LAT-1 Lectin Domain |
|---|---|
| **Data Collection** | |
| Beamline | APS 24-ID-E |
| Space Group | $P2_1$ |
| *Cell Dimensions* | |
| $a, b, c$ (Å) | 74.152, 316.755, 172.441 |
| $\alpha, \beta, \gamma$ (°) | 90, 90, 90 |
| Resolution (Å) | 200-4.00 (4.42-4.00)* |
| $R_{sym}$ (%) | 11.2 (72.7) |
| $\langle I \rangle / \langle \sigma(I) \rangle$ | 6.2 (2.1) |
| $CC_{1/2}$ | 0.996 (0.603) |
| Completeness (%) (spherical) | 55.6 (16.7) |
| (ellipsoidal)† | 88.3 (80.2) |
| Redundancy | 3.4 (3.3) |
| **Refinement** | |
| Resolution (Å) | 74.15-4.00 (4.11-4.00)* |
| Reflections | 37,091 |
| $R_{cryst}$ (%) | 29.77 (31.54) |
| $R_{free}$ (%)** | 33.73 (44.13) |
| *Number of atoms* | |
| Protein | 32,410 |
| Ligand/Glycans | 530 |
| Water | 0 |
| *Average B-factors (Å$^2$)* | |
| All | 194.2 |
| Protein | 194.8 |
| Ligand/Glycans | 161.1 |
| Solvent | N/A |
| *R.m.s. deviations from ideality* | |
| Bond Lengths (Å) | 0.004 |
| Bond Angles (°) | 0.890 |
| *Ramachandran plot* | |
| Favored (%) | 86.68 |
| Outliers (%) | 0.31 |
| Rotamer Outliers (%) | 1.48 |
| All-atom Clashscore‡ | 14.22 |

* The values in parentheses are for reflections in the highest resolution bin.

** 5% of reflections (1,858) was not used during refinement for cross validation purposes.

† Diffraction limits (Å) and corresponding principal axes of the ellipsoid fitted to the diffraction cut-off surface as direction cosines in the orthogonal basis (standard PDB convention), and in terms of reciprocal unit-cell vectors, are:

| | | | |
|---|---|---|---|
| Diffraction limit #1: | 6.138 Å | (0.9749, 0.0000, –0.2226) | 0.881 **a*** – 0.473 **c*** |
| Diffraction limit #2: | 3.548 Å | (0.0000, 1.0000, 0.0000) | **b*** |
| Diffraction limit #3: | 4.534 Å | (0.2226, 0.0000, 0.9749) | 0.098 **a*** + 0.995 **c*** |

Eigenvalues of overall anisotropy tensor on |F|s (Å$^2$) and corresponding eigenvectors of the overall anisotropy tensor as direction cosines in the orthogonal basis (standard PDB convention), and in terms of reciprocal unit-cell vectors:

| | | | |
|---|---|---|---|
| Eigenvalue #1: | 306.72 | (0.9993, 0.0000, –0.0371) | 0.996 **a*** – 0.093 **c*** |
| Eigenvalue #2: | 121.51 | (0.0000, 1.0000, 0.0000) | **b*** |
| Eigenvalue #3: | 213.72 | (0.0371, 0.0000, 0.9993) | 0.016 **a*** + c* |

‡ Clashscores were calculated by *phenix.refine* (*Phenix* version 1.20.1).

N/A: Not applicable.
